# Pyrrolocytosine as a Fluorescent Sensor of RNA-Small Molecule Interactions

**DOI:** 10.64898/2026.09.29.755477

**Authors:** Taylor L. Coulson, Gavin Valdez, Julia R. Widom

## Abstract

Fluorescent base analogues have proven to be valuable probes of RNA structure. Pyrrolocytosine (pC), an analogue of cytosine, exhibits highly environment-sensitive fluorescence within RNA and is strongly quenched in hydrogen-bonded contexts. In this work, we incorporated pC into the aptamer domain of the B. subtilis guanine riboswitch to determine if this quenching mechanism could allow for sensing of RNA- small molecule interactions. We utilized complementary fluorescence spectroscopy methods to understand the functional impacts and ligand-sensitivity of pC substitutions distal to and within the binding pocket. We found that pC substitution within a static structural region exhibits the expected ligand-independent fluorescence properties. When placed in the ligand binding pocket, pC exhibits a moderately decreased affinity for the ligands guanine (G) and hypoxanthine (Hx) while maintaining high specificity for G. We found that pC behaves as a turn-off sensor, with distinct changes in fluorescence upon G or Hx binding. These differences are dependent on the number of hydrogen bonds formed with pC and are resolvable in both time-dependent and steady-state fluorescence measurements. Our results show that pC can be used to study RNA-small molecule interactions and can discriminate between similar ligands even when they have identical binding modes.

## 1. INTRODUCTION

Decades of RNA research have expanded our understanding of its biological roles from a simple carrier of genetic information to a molecule with wide-ranging regulatory capacity. Regulatory functions are commonly achieved by non-coding RNAs (ncRNAs), which are transcribed from regions of the genome that do not encode protein, as well as non-coding domains of messenger RNAs (mRNAs) termed “untranslated regions” (UTRs). These transcripts and motifs have been shown to be required for regulation of gene expression in bacteria [1] and their dysregulation has been identified in multiple disease states [2]. As a result, ncRNAs and motifs found in UTRs have seen increased research interest for their potential as therapeutic targets [3–6]. Riboswitches, which are found primarily in the 5’ UTRs of bacterial mRNAs, represent one class of RNA motifs that has garnered such interest. Riboswitches control expression of downstream genes in response to environmental stimuli, typically at the transcriptional or translational level. These molecular switches consist of two domains, the aptamer domain and the expression platform. The aptamer domain binds ligands in the cellular environment and undergoes a conformational change upon binding; this induces a further structural change in the expression platform through a switching sequence that links the two domains, resulting in ligand-dependent regulation of the downstream genes [7,8]. Structure-guided modification of native riboswitch ligands has been shown to be a promising avenue of initial drug candidate design [9,10], and novel synthetic ligands targeting riboswitches have shown regulatory effects in different bacterial strains [11–13]. A more thorough understanding of how ligand binding is coupled to the local conformation within the aptamer domain can aid in the design of novel ligands that may have potential as drug candidates.

Fluorescence-based methods are widely used to investigate RNA’s structure in solution and how it changes in response to environmental stimuli, though this approach requires labeling with a minimally perturbing probe that is sensitive to the RNA’s structure [14–17]. For example, fluorescent base analogues (FBAs) are structural mimics of native nucleic acid bases with increased fluorescence quantum yields and can be incorporated into an RNA at a location of interest [18,19]. FBAs are typically designed to maintain the base pairing of their corresponding native bases, though the structural modifications implemented for improved fluorescence have the potential to perturb RNA tertiary structure [20]. Many FBAs exhibit environment-sensitive fluorescence [21–24], making them useful probes of local nucleic acid structure, and new FBAs with distinct photophysical and structural properties are constantly being developed [25–28].

Pyrrolocytosine (pC; Fig. 1A) was first utilized as a deoxycytidine analogue to in-vestigate formation of the RNA:DNA hybrid in the transcription bubble [29]. More recent investigations have utilized pC to investigate folding of a ribozyme [30] and intermolecular interactions between tRNA and the T-Box riboswitch [31]. Previous experimental and computational studies showed that within DNA oligonucleotides, pC exhibits fluorescence enhancement in base-stacked conformations while hydrogen bonding quenches its fluorescence [32,33]. Building on this work, we recently showed that while the established photophysical impacts of base stacking and hydrogen bonding persist in RNA, there is also a strong sequence dependence not previously reported in DNA. Additionally, we showed that the base-paired state of pC within a duplex can be resolved using fluorescence lifetime measurements as a highly populated, rapidly decaying time component, and that the lineshape of its emission spectrum is a reporter of local polarity [34]. These results supported the notion that pC could act as a multifaceted probe of local structure when utilized in more complex RNA species.

**Figure 1.**
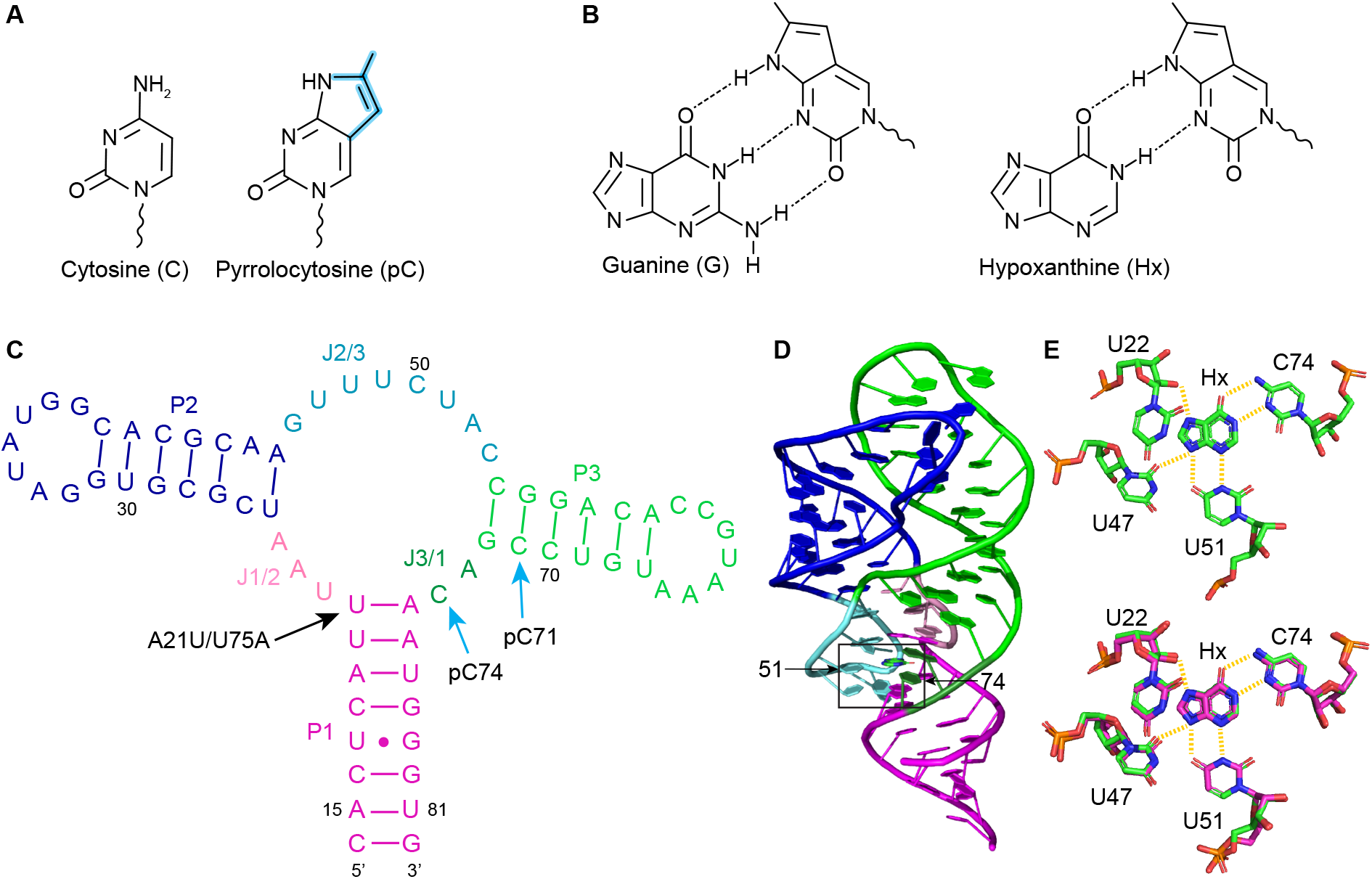
Pyrrolocytosine and the guanine riboswitch. (**A**) Chemical structures of cytosine (left) and pyrrolocytosine (right). Modifications are highlighted in blue on the pC structure. (**B**) Hydrogen bonding patterns between pC and guanine (left) or hypoxanthine (right). (**C**) Sequence and secondary structure of the A21U/U75A mutant of the B. subtilis GR utilized in this study. Sites of pC labeling are indicated. (**D**) Crystal structure of the Hx-bound A21U/U75A mutant of the B. subtilis GR (PDB ID 2EES) with domains color-coded as in panel C. (**E**) Top: Close-up of the binding pocket (box in panel D) showing hydrogen bonds between Hx and nearby bases. Bottom: Overlay of binding pocket structures of WT (PDB ID 4FE5) and A21U/U75A (PDB ID 2EES) GR variants. Structures are color-coded by atom, where C is green (2EES) or pink (4FE5).

Most studies that utilize FBAs, including pC, do so by labeling sites that undergo environment-sensitive reorganization of stacking or base-pairing interactions [30,31,35]. A yet unexplored application of pC’s unique fluorescence properties is the investigation of small molecule interactions with RNA. While the widely used 2-aminopurine (2-AP) has been used to understand the ligand-dependent conformational changes of riboswitches, it has typically been placed in distal regions of the aptamer domain that do not directly interact with the ligand [36–39] or in rarer cases has been used as a ligand analogue [40–42].

While 2-AP and many other FBAs are quenched primarily through base stacking [43,44], pC’s hydrogen-bond-dependent quenching presents the opportunity for its application as a direct turn-off sensor of ligand binding in riboswitches that rely on hydrogen bonds between the ligand and a cytosine residue (Fig. 1B). The guanine riboswitch (GR; Fig. 1C-D) aptamer domain coordinates guanine (G) and its metabolic derivatives with varying affinity through a Watson-Crick-Franklin (WCF) hydrogen bonding pattern to a conserved cytosine base, C74 (Fig. 1E) [41,45]. This riboswitch acts through a hybrid conformational selection/induced fit model [41]; its three-way junction is largely pre-organized while an unhybridized linker (J2/3, Fig. 1C) between two adjacent stem-loops samples “open” and “closed” conformations, where only the open conformation is capable of binding ligand [46,47]. We hypothesized that pC substituted for C74 could act as a conformationally sensitive reporter of ligand binding, while pC substituted elsewhere could reveal whether ligand binding is coupled to conformational changes distal to the binding pocket.

In this work, we utilized both steady-state and time-dependent spectroscopic methods to monitor pC’s fluorescence properties within GR in response to the binding of ligands with varying affinities and degrees of hydrogen bonding (Fig. 1B). pC substituted within the P3 helix confirmed that this domain is base-paired both with and without ligand, while pC substituted for C74 exhibited ligand-dependent quenching while maintaining specificity, as assessed through both equilibrium binding affinity and binding kinetics. The emission lineshape of pC indicated different levels of solvent accessibility at these two labeling sites, and both sites showed evidence of conformational heterogeneity both with and without ligand. These results demonstrate that pC can be a valuable tool for measuring the conformational states accessible to riboswitches using both steady-state and time-dependent methods.

## 2. RESULTS

### 2.1. Design of pC-based GR sensors

The GR was selected as a model system for this study as its ligand specificity is conferred by a key cytosine residue [41,48] (C74) and it has longstanding interest as a pharmaceutical target [10–13,49]. Specifically, the *Bacillus subtilis* GR aptamer domain has been the subject of extensive biochemical and structural characterization [41,45,47,50–54], allowing for informed labeling site selection and guiding interpretation of experimental results. We selected two pC labeling sites within a 69-nucleotide sequence encompassing the *B. subtilis* GR aptamer domain: positions 71 and 74 (Table 1). Position 71 resides within P3 (Fig. 1C) and remains base-paired independently of Mg^2+^ and ligand [52]. pC71 is therefore expected to remain in a highly quenched state, defined by a low emission intensity and short fluorescence lifetime, and observations to the contrary would be indicative of pC altering the surrounding structure.

**Table 1.**
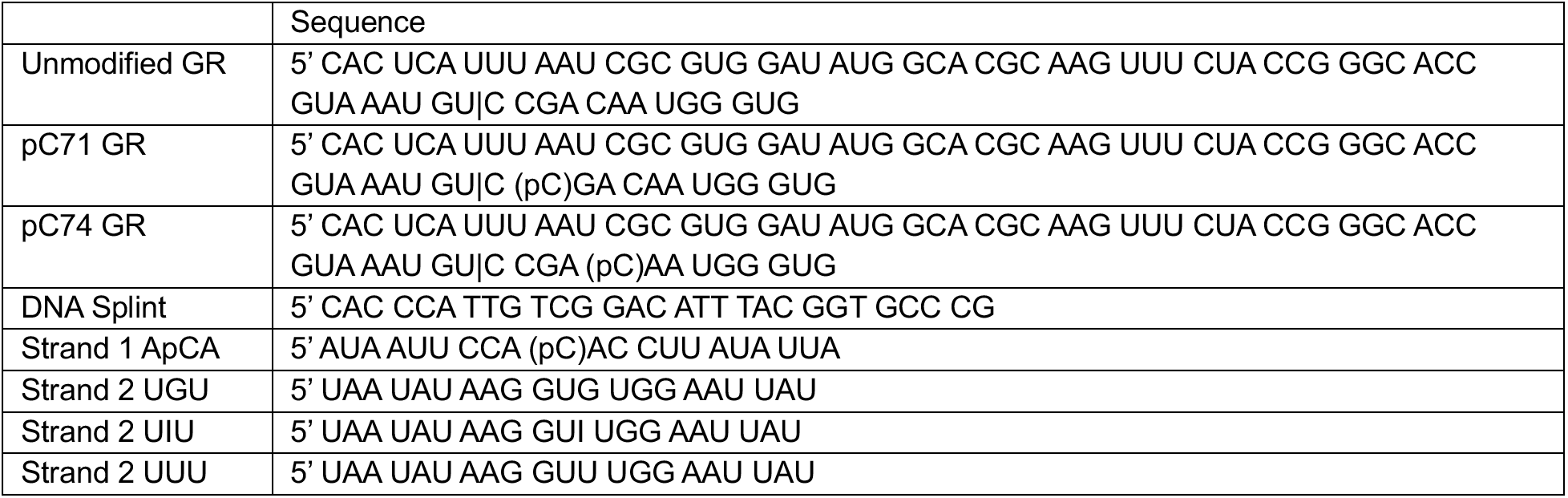
Sequences of oligonucleotides used in this study. A vertical line indicates the site at which two oligonucleotides were ligated together to form the full-length GR.

The binding affinity and specificity of GR is determined primarily by the extent of hydrogen bonding between a potential ligand and C74 [41,45,48]. pC74 is therefore expected to experience strong fluorescence quenching in the presence of ligand, as we have previously demonstrated that base pairing of pC with G results in reduced emission intensity and a dominant fast decay component [34]. In order to test whether pC74 GR maintained the ligand binding properties of the native GR and to investigate the dependence of pC fluorescence on the number of hydrogen bonds formed, we carried out all measurements with both G, which forms three hydrogen bonds with C, and a lower affinity ligand, hypoxanthine (Hx), which lacks a C2 primary amine and therefore only forms two hydrogen bonds with C74 (Fig. 1B).

To optimize this sensor, the sequence context of pC also had to be considered. We previously demonstrated that pC’s fluorescence within RNA is strongly sequence-dependent, with adjacent A and C residues conferring the greatest fluorescence enhancement over free pC riboside. Adjacent U residues confer the weakest enhancement and yield three-component fluorescence decays, leading to a more complex structural interpretation than required in other sequence contexts [34]. In the selected *B. subtilis* GR, C74 resides between A73 and U75, the latter of which makes a base-pair with A21. Based on our previous results showing divergent properties of pC when adjacent to A and U, we asked whether the sequence might offer any flexibility that could be exploited to improve performance. Interestingly, nucleotides 21 and 75 have been shown to covary across species to maintain the A-U WCF base pair (A21/U75 in 90% of species and U21/A75 in 10%) [55] and mutational studies have shown that affinity for G is identical in WT and A21U/U75A and is impaired only slightly in A21G/U75C [53,56]. The affinity for Hx is affected, with a KD of 0.76 µM for WT GR and 3.3 µM for A21U/U75A at 30 °C in 10 mM Mg^2+^ [56]. However, x-ray crystallography of GR bound to Hx has shown that its structure is unchanged by the A21U/U75A transversion, including the hydrogen bonding network around the ligand (Fig. 1E), and the adjacent “platform” created by the 21-75 base pair and C50 [53]. We therefore hypothesized that a pC-labeled A21U/U75A mutant could act as a functional sensor.

A common quenching mechanism for FBAs is the formation of low-energy charge-transfer (CT) states [57–61], so we performed quantum mechanical (QM) calculations to ask whether they were likely to be present in any of the sequence contexts under consideration. QM calculations performed on the A-form trimers ApCA, UpCU and ApCU all showed an absence of low-energy CT states, with S0-S1 being a strong π-π* transition localized on pC (Fig. S1), similar to our previous observations with CpCC and GpCG. The S2 state was at least 0.6 eV higher in energy in all cases. The same observations held when QM calculations were performed on ApCA and ApCU trimers in the geometry of the GR binding pocket observed crystallographically (PDB: 4FE5 [45], 2EES [53]; Fig. S1). While these results did not reveal an obvious photophysical justification for any particular sequence, it remained advantageous to obtain maximal pC fluorescence intensity and more readily interpretable results by removing sequence-dependent variables where possible. We therefore placed pC74 in the more favorable ApCA context by introducing the compensatory mutation A21U/U75A (Fig. 1C), taking advantage of the biochemical and structural characterization discussed above.

### 2.2. pC71 exhibits stable base pairing

We first consider pC71, which replaces a base-paired C residue within the P3 helix (Fig. 1C). This helix is intact in the isolated RNA [52], so we anticipated a low emission intensity and little change in the fluorescence properties upon addition of Mg^2+^ or ligand. Indeed, emission and excitation spectra of pC71 GR show no change in intensity or lineshape upon addition of 1 mM Mg^2+^, and only a small decrease in intensity upon addition of saturating G or Hx (4 µM or 310 µM, respectively) (Fig. 2A-B). The initial low emission intensity suggests that pC71 is in a base-paired state, and the observed decrease upon addition of G/Hx could be attributed to increased base-pairing stability within P3. As C71 does not directly interact with the bound ligand, this consistent response indicates that pC71 does not disrupt the global structure of the riboswitch. Of particular note is the distinct emission lineshape pC71 GR exhibits relative to previously reported pC-labeled RNAs. While the suppressed “red shoulder” associated with highly stacked and solvent shielded environments [34] is clearly visible at ∼470 nm, there is an additional longer wavelength (∼530 nm) feature indicating a distinct solvation environment within this highly ordered region of GR (Fig. 2A; Fig. S2A).

**Figure 2.**
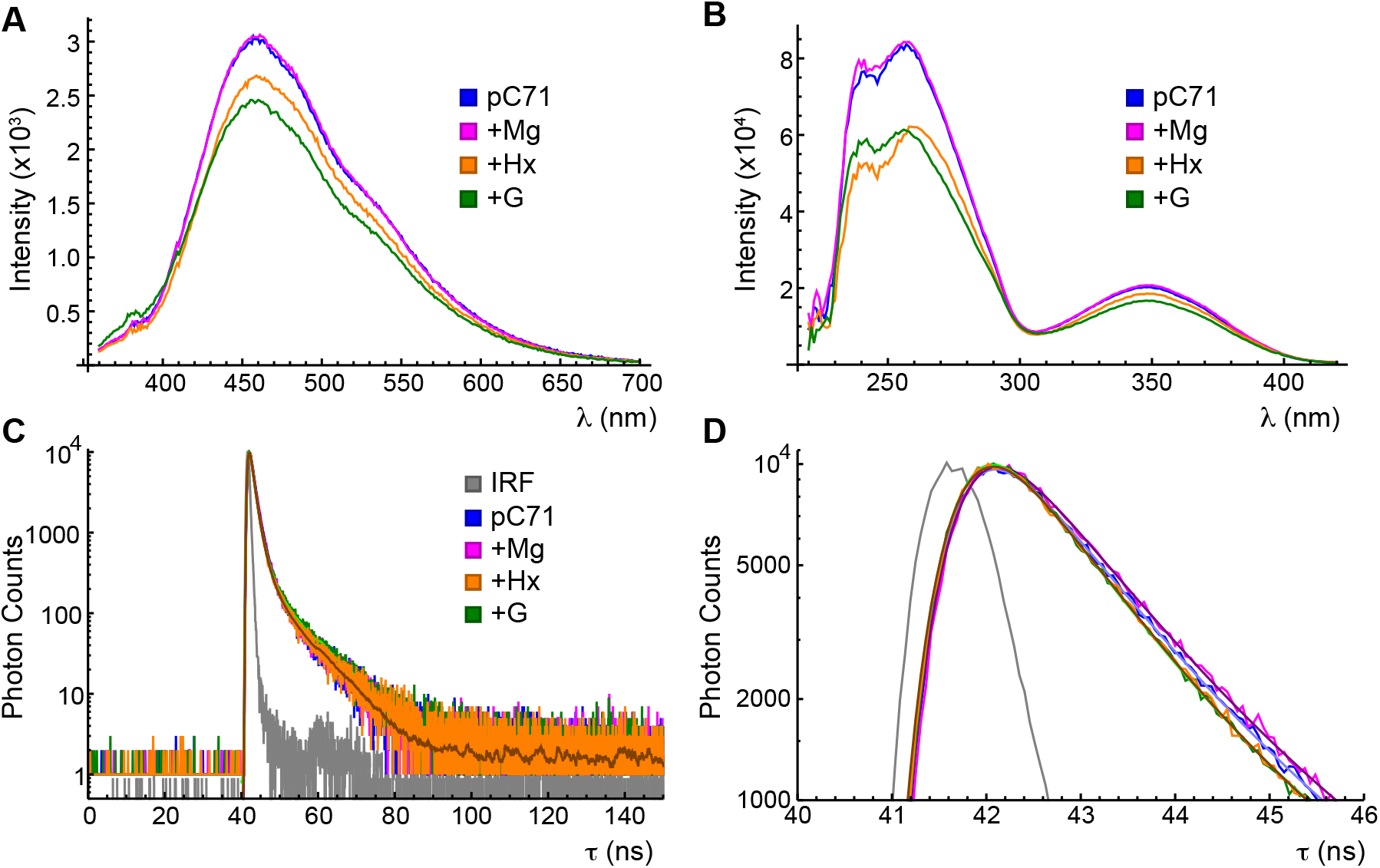
Steady-state and time-resolved spectroscopy of pC71 GR. (**A**) Fluorescence emission spectra of pC71 GR in standard phosphate buffer (blue), buffer supplemented with 1 mM MgCl_2_ (magenta), buffer supplemented with 1 mM MgCl_2_ and 310 µM Hx (orange) or buffer supple-mented with 1 mM MgCl_2_ and 4 µM G (green). (**B**) Fluorescence excitation spectra of pC71 GR. Color coding as in panel A. (**C**) Fluorescence decays of pC71 GR. Color coding as in panel A, with a typical instrument response function (IRF) additionally plotted in gray. 1E1G_S_ fits are overlaid in contrasting shade. (**D**) Close-up of the short-time region of panel C.

The hydrogen-bonded state of pC71 is further confirmed by time-correlated single photon counting (TCSPC; Fig. 2C-D). In these measurements, all emission at wavelengths longer than 420 nm was collected (see Methods 3.3.2); thus, all spectral features discussed above are represented in the resulting data. pC71 fluorescence decays were well fit (χ^2^=1.10) with a model consisting of a slow exponential decay component and a shorter component described by a gamma distribution [62], termed a “1E1G_S_” model (details in Appendix A). This model returned lifetimes of 1.1 and 8.4 ns with normalized amplitudes (termed “population weights”) of 0.99 and 0.01, respectively (Table 2). The population weight estimates the occupancy of a particular conformation, or a set of rapidly interconverting conformations, of pC relative to the others that are temporally resolvable, under the assumption that the conformations have equal excitation and detection efficiencies and radiative decay rates to one another. Decay components of 0.5-3 ns have been consistently observed in a variety of base-paired sequence contexts [31,34], so the dominance of the 1.1 ns component indicates that pC71 is constrained to a primarily base-paired conformation within P3. The 8.4 ns component likely arises from rare breathing events resulting in a base-stacked but un-paired conformation. No significant change in the lifetime or population weight of either component is observed when G or Hx is added. The ligand-independent emission behavior of pC71 GR, coupled with its circular dichroism (CD) spectrum being unchanged relative to the unmodified GR (Fig. S3), indicates that pC substitution minimally perturbs highly structured regions within the RNA.

**Table 2.** TCSPC fitting results for pC71 GR. Uncertainties represent the standard deviation across 3 successive data acquisitions. Unless otherwise stated, all data was collected at pH 7.4.

| pC71 Condition | Model | $\chi^2$ | $\langle\tau\rangle_{\text{obs}}$ (ns) | $w_1$ | $\tau_1$ (ns) | $w_2$ | $\tau_2$ (ns) | $w_3$ | $\tau_3$ (ns) |
| --- | --- | --- | --- | --- | --- | --- | --- | --- | --- |
| Alone | 1E1Gs | 0.95 | 1.190<br>$\pm 0.005$ | 0.0127<br>$\pm 0.0009$ | 8.4<br>$\pm 0.3$ | 0.9873<br>$\pm 0.0009$<sup>1</sup> | 1.097<br>$\pm 0.004$<sup>1</sup> | -- | -- |
| +1 mM Mg<sup>2+</sup> | 1E1Gs | 0.96 | 1.24<br>$\pm 0.03$ | 0.0125<br>$\pm 0.0005$ | 8.5<br>$\pm 0.2$ | 0.9875<br>$\pm 0.0005$<sup>1</sup> | 1.15<br>$\pm 0.03$<sup>1</sup> | -- | -- |
| +4 μM G | 1E1Gs | 1.03 | 1.08<br>$\pm 0.01$ | 0.014<br>$\pm 0.001$ | 8.6<br>$\pm 0.2$ | 0.986<br>$\pm 0.001$<sup>1</sup> | 0.968<br>$\pm 0.006$<sup>1</sup> | -- | -- |
| +310 μM Hx | 1E1Gs | 0.96 | 1.09<br>$\pm 0.01$ | 0.0134<br>$\pm 0.0004$ | 8.0<br>$\pm 0.1$ | 0.9865<br>$\pm 0.0004$<sup>1</sup> | 1.00<br>$\pm 0.01$<sup>1</sup> | -- | -- |
| Alone, pH 4 | 3E | 1.28 | 2.3<br>$\pm 0.01$ | 0.044<br>$\pm 0.003$ | 9.1<br>$\pm 0.1$ | 0.384<br>$\pm 0.004$ | 3.52<br>$\pm 0.05$ | 0.572<br>$\pm 0.006$ | 0.97<br>$\pm 0.01$ |

<sup>1</sup> Parameter originates from a gamma component.

### 2.3. Model RNAs reveal the photophysical impacts of hydrogen bonding between pC and G or Hx

To build a foundation for understanding the behavior of pC74 in the complex environment of the binding pocket, we investigated model 21-nucleotide single-stranded (ss) and double-stranded (ds) RNAs containing a central ApCA motif (Table 1; Fig. 3). Model dsRNAs contained a central ApCA:UGU motif (termed 1:2UGU) to mimic the guanine-bound riboswitch, an ApCA:UIU motif (1:2UIU; I refers to inosine, the nucleoside with Hx as its base) to mimic the Hx-bound riboswitch, or a mismatched ApCA:UUU motif (1:2UUU). TCSPC measurements were performed and, based on reduced χ^2^ values and the behavior of the residuals (details in Appendix A), a 2E model was selected to describe the decays of the pC-labeled ssRNA St1ApCA and a 3E model was selected to describe the decays of 1:2UGU, 1:2UIU and 1:2UUU (Table 3). Under the established structural interpretation of pC fluorescence decays [34], St1ApCA exhibits a dominant (population weight of 0.7) base-stacked conformation with a lifetime of 11.9 ns and a less populated unstacked conformation with lifetime 4.6 ns.

**Figure 3.**
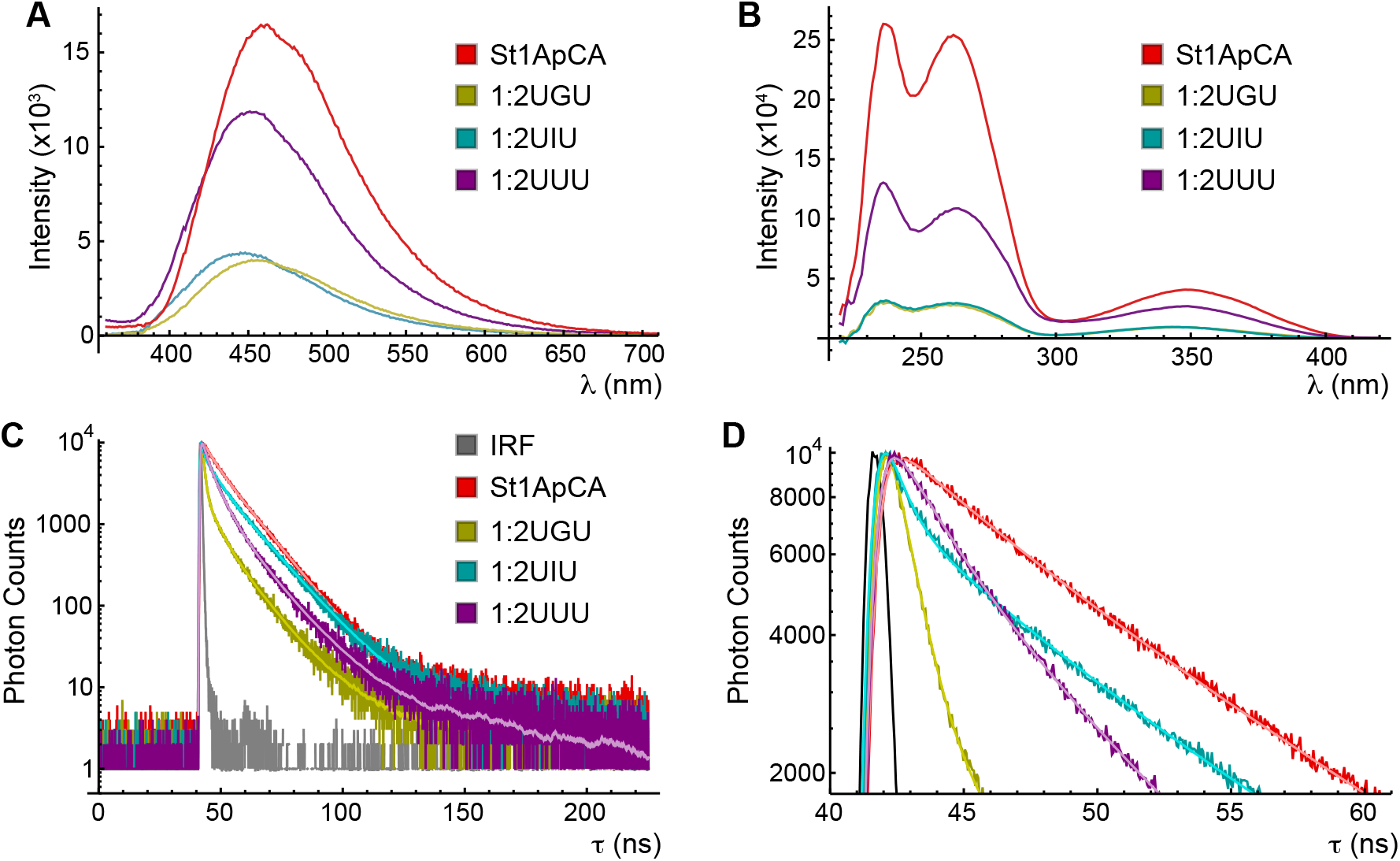
Steady-state and time-resolved spectroscopy of model ssRNA and dsRNAs. (**A**) Fluorescence emission spectra of St1ApCA (red), 1:2UGU (yellow), 1:2UIU (cyan) and 1:2UUU (purple). (**B**) Fluorescence excitation spectra of model RNAs. Color coding as in panel A. (**C**) Fluorescence decays of model RNAs. Color coding as in panel A, with a typical IRF additionally plotted in gray. 2E (for St1ApCA) or 3E (for all others) fits are overlaid in contrasting shade. (**D**) Close-up of the short-time region of panel C.

**Table 3.** TCSPC fitting results for pC riboside (pCr) and model 21-mers. Uncertainties represent the standard deviation across 3 successive data acquisitions. Unless otherwise stated, all data was collected at pH 7.4.

| RNA | Model | $\chi^2$ | $\langle\tau\rangle_{\text{obs}}$<br>(ns) | $w_1$ | $\tau_1$<br>(ns) | $w_2$ | $\tau_2$<br>(ns) | $w_3$ | $\tau_3$<br>(ns) |
| --- | --- | --- | --- | --- | --- | --- | --- | --- | --- |
| pCr | 2E | 1.11 | 1.904<br>$\pm 0.005$ | 0.015<br>$\pm 0.008$ | 5.1<br>$\pm 0.8$ | 0.985<br>$\pm 0.008$ | 1.86<br>$\pm 0.01$ | -- | -- |
| pCr, pH 4 | 2E | 1.07 | 1.878 | 0.0073 | 7.2 | 0.9927 | 1.839 | -- | -- |
| | | | $\pm 0.003$ | $\pm 0.0008$ | $\pm 0.4$ | $\pm 0.0008$ | $\pm 0.004$ | | |
| St1ApCA | 2E | 1.27 | 9.71<br>$\pm 0.02$ | 0.701<br>$\pm 0.002$ | 11.90<br>$\pm 0.02$ | 0.299<br>$\pm 0.002$ | 4.56<br>$\pm 0.08$ | -- | -- |
| 1:2UGU | 3E | 1.12 | 1.83<br>$\pm 0.02$ | 0.037<br>$\pm 0.003$ | 13.4<br>$\pm 0.3$ | 0.1016<br>$\pm 0.0006$ | 5.6<br>$\pm 0.3$ | 0.862<br>$\pm 0.003$ | 0.90<br>$\pm 0.02$ |
| 1:2UIU | 3E | 1.28 | 4.6<br>$\pm 0.2$ | 0.230<br>$\pm 0.008$ | 13.35<br>$\pm 0.08$ | 0.193<br>$\pm 0.007$ | 5.5<br>$\pm 0.1$ | 0.58<br>$\pm 0.01$ | 0.57<br>$\pm 0.04$ |
| 1:2UUU | 3E | 1.18 | 5.33<br>$\pm 0.01$ | 0.138<br>$\pm 0.002$ | 13.49<br>$\pm 0.08$ | 0.51<br>$\pm 0.01$ | 5.44<br>$\pm 0.03$ | 0.35<br>$\pm 0.01$ | 1.99<br>$\pm 0.06$ |
| St1ApCA,<br>pH 4 | 3E | 1.22 | 3.168<br>$\pm 0.007$ | 0.046<br>$\pm 0.007$ | 11.5<br>$\pm 0.5$ | 0.40<br>$\pm 0.01$ | 4.6<br>$\pm 0.1$ | 0.56<br>$\pm 0.02$ | 1.45<br>$\pm 0.05$ |

3E fits to the fully complementary duplexes 1:2UGU and 1:2UIU each revealed a sub-nanosecond decay component with lifetime of 0.90 or 0.57 ns and population weight of 0.86 or 0.58, respectively. These sub-ns lifetimes are within the practical lower limit of detection of the instrument (see Methods 3.3.2), and have been previously reported for pC in measurements of similar time resolution [31]. We assign this component to a base-paired conformation, analogous to the dominant 0.5-3 ns components previously observed in dsRNA across several sequence contexts [31,34] and the dominant 1.1 ns component observed in pC71 GR (Table 2). All three duplexes exhibit a ∼5.5 ns component with a population weight inversely related to the degree of pC hydrogen bonding: 0.10 with pC opposite G, 0.19 with pC opposite I, and 0.51 with pC opposite U. We assign this component to an un-stacked conformation, analogous to the 4.4 ns component observed in St1ApCA. All three duplexes additionally exhibit a ∼13 ns component that was least populated with pC opposite G (w=0.04) and most populated with pC opposite I (w=0.23). We assign this to a “breathed open” conformation in which pC is stacked but not base-paired, analogous to the 12 ns component observed in St1ApCA. Interestingly, the mismatched duplex 1:2UUU also exhibits a 2 ns decay component (w=0.35), which was not previously observed with a CpCC:GUG motif [34] (discussed further below). In summary, the fluorescence decays of the dsRNAs investigated here reveal stacked and unstacked populations analogous to those observed in other model RNAs, as well as a faster decay of ≤2 ns whose population weight is related to the degree of hydrogen bonding between pC and the opposing base (opposite G > opposite I > opposite U).

### 2.4. pC74 exhibits ligand-dependent fluorescence quenching and maintains binding specificity

We next examine pC74, which replaces the ligand-coordinating C in the J3/1 linker (Fig. 1C). As pC retains the necessary hydrogen-bonding pattern (Fig. 1B), we anticipated that pC74 GR would exhibit large ligand-dependent changes in its fluorescence properties. pC74 GR exhibits the expected fluorescence enhancement of pC in base-stacked conformations, resulting in a similar brightness as observed in St1ApCA (Fig. 3A, 4A). In the presence of ligand, the emission spectra of pC74 GR distinguish between G and Hx, where the reduction in overall intensity is directly related to the number of hydrogen bonds formed with the ligand (Fig. 4A). The addition of a saturating concentration of G results in an ∼40% decrease in intensity while Hx only induces an ∼30% reduction. The emission lineshape of pC74 exhibits the suppressed red shoulder at ∼470 nm that we have shown is indicative of a base-stacked or non-polar environment. There is no discernable change in the lineshape of the emission spectra in either ligand-bound complex (Fig. S2C), in contrast to the blueshift and further suppression of the red shoulder observed upon hybridization of pC-labeled ssRNA into dsRNA (Fig. S2E) [34].

**Figure 4.**
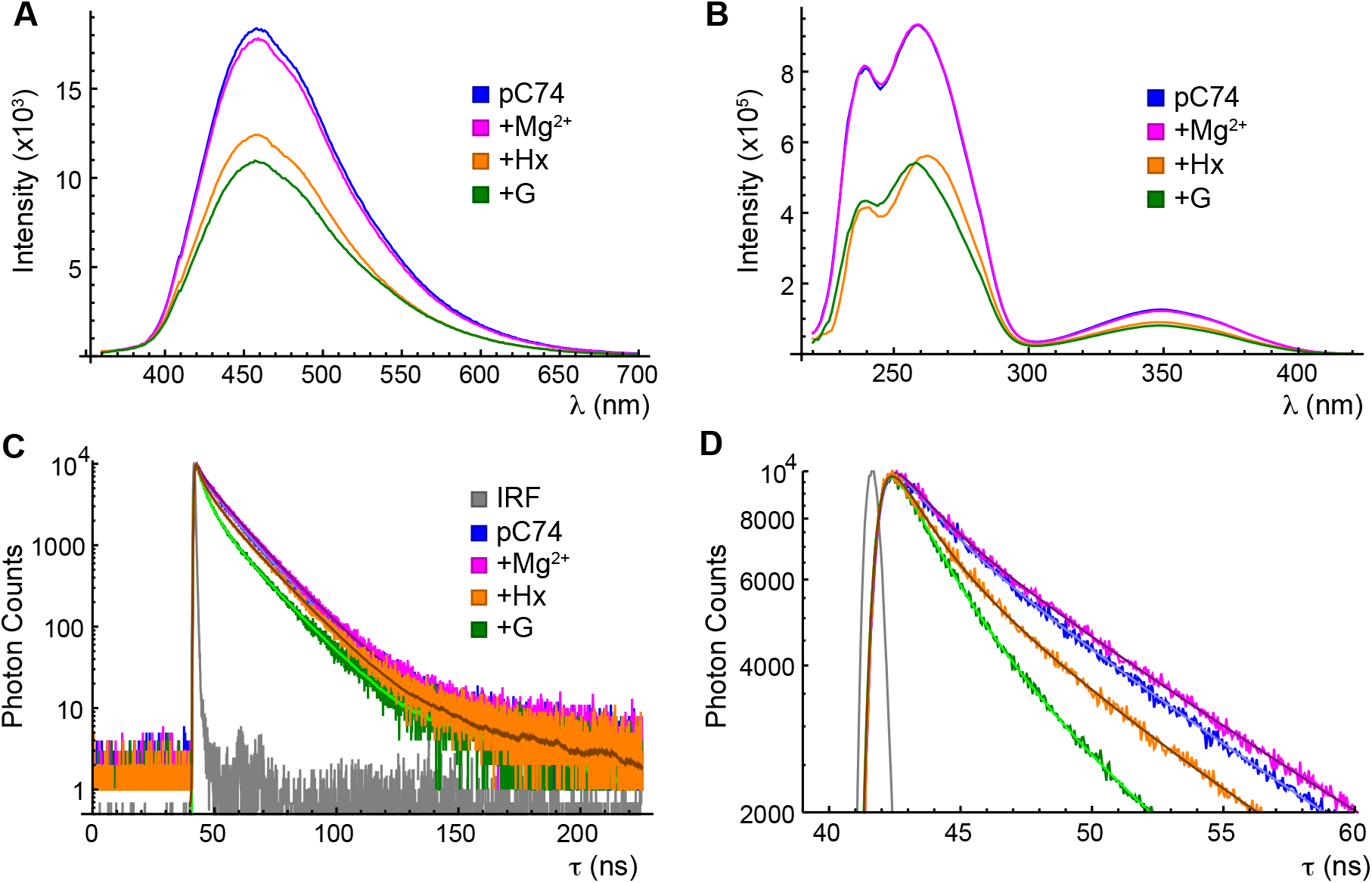
Steady-state and time-resolved spectroscopy of pC74 GR. (**A**) Fluorescence emission spectra of pC74 GR in standard phosphate buffer (blue), buffer supplemented with 1 mM MgCl_2_ (magenta), buffer supplemented with 1 mM MgCl_2_ and 310 µM Hx (orange) or buffer supplemented with 1 mM MgCl_2_ and 4 µM G (green). (**B**) Fluorescence excitation spectra of pC74 GR. Color coding as in panel A. (**C**) Fluorescence decays of pC74 GR. Color coding as in panel A, with a typical IRF additionally plotted in gray. 3E fits are overlaid in contrasting shade. (**D**) Close-up of the short-time region of panel C.

We have previously demonstrated that the fluorescence excitation spectrum of pC exhibits signatures of the surrounding sequence context due to energy transfer from nearby native bases, which yields a distinct spectral feature around 260 nm [34]. Upon the addition of G to pC74 GR, the peak narrows with a reduction in intensity on the red edge (Fig. S2D). This change, though slight, could be indicative of reduced interaction with the adjacent A residues (λ_max_ = 259 nm for adenosine monophosphate [63]) or direct energy transfer from the hydrogen-bonded G (λ_max_ = 246 nm; Fig. S4), which we have previously observed in dsRNA [34]. No change is observed in this feature upon addition of G to pC71 GR (Fig. S2B), supporting the notion that the subtle shift observed in pC74 GR arises from altered geometry within the binding pocket. As indicated by excitation spectra of the model 21-mers (Fig. S2F), shifts in energy transfer interactions between pC and adjacent residues are more subtle in an ApCA context than previously observed in a CpCC context [34]. This is likely due to the smaller difference between the λ_max_ of the A residues flanking pC and those of G and U residues opposite it in the duplexes studied here (λ_max_ = 252 nm for guanosine monophosphate, λ_max_ = 262 nm for uridine monophosphate).

While the excitation spectrum of pC74 GR in the presence of Hx does exhibit a clear change in the λ_max_ of this peak to 262 nm (Fig. 4B; Fig. S2), this is likely confounded by the large absorbance of the saturating Hx and therefore larger inner filter effect correction required (Fig. S4). A similar shift in the fluorescence excitation spectrum is observed upon addition of Hx to pC71 GR (Fig. S2B), confirming that it does not reflect alterations in the binding pocket.

Exploiting the fluorescence quenching exhibited by pC74 in the presence of ligand, fluorescence titrations were carried out in order to determine the K_D_ values for G and Hx at physiological (1 mM) and high (20 mM) Mg^2+^ concentrations for comparison to literature (Fig. 5, Fig. S5). In 20 mM Mg^2+^, a 4- parameter logistic (4PL) fit to maximum emission intensity vs. total ligand concentration yielded apparent K_D_ values 35±3 nM for G and 12.1±0.2 µM for Hx (mean ± standard deviation across three experimental replicates). K_D_ values for the A21U/U75A mutant of the GR were previously determined via isothermal titration calorimetry in 10 mM Mg^2+^, and were found to be 3 nM for G and 3.3 µM for Hx [56]. The authors noted no change in the binding affinity at higher concentrations of Mg^2+^. In 1 mM Mg^2+^, pC74 exhibited apparent K_D_ values of 80±10 nM for G and 41±4 µM for Hx. Substitution of pC at position 74 in an A21U/U75A background thus modestly decreases the binding affinity while maintaining apparent K_D_ values of the same order of magnitude and high specificity for G over Hx. It should be noted that the actual K_D_ values for G are likely lower than the K_app_ values obtained from the 4PL fits, as significant depletion of G is likely to occur with the 150 nM RNA concentration utilized here (see Methods 3.3.1).

**Figure 5.**
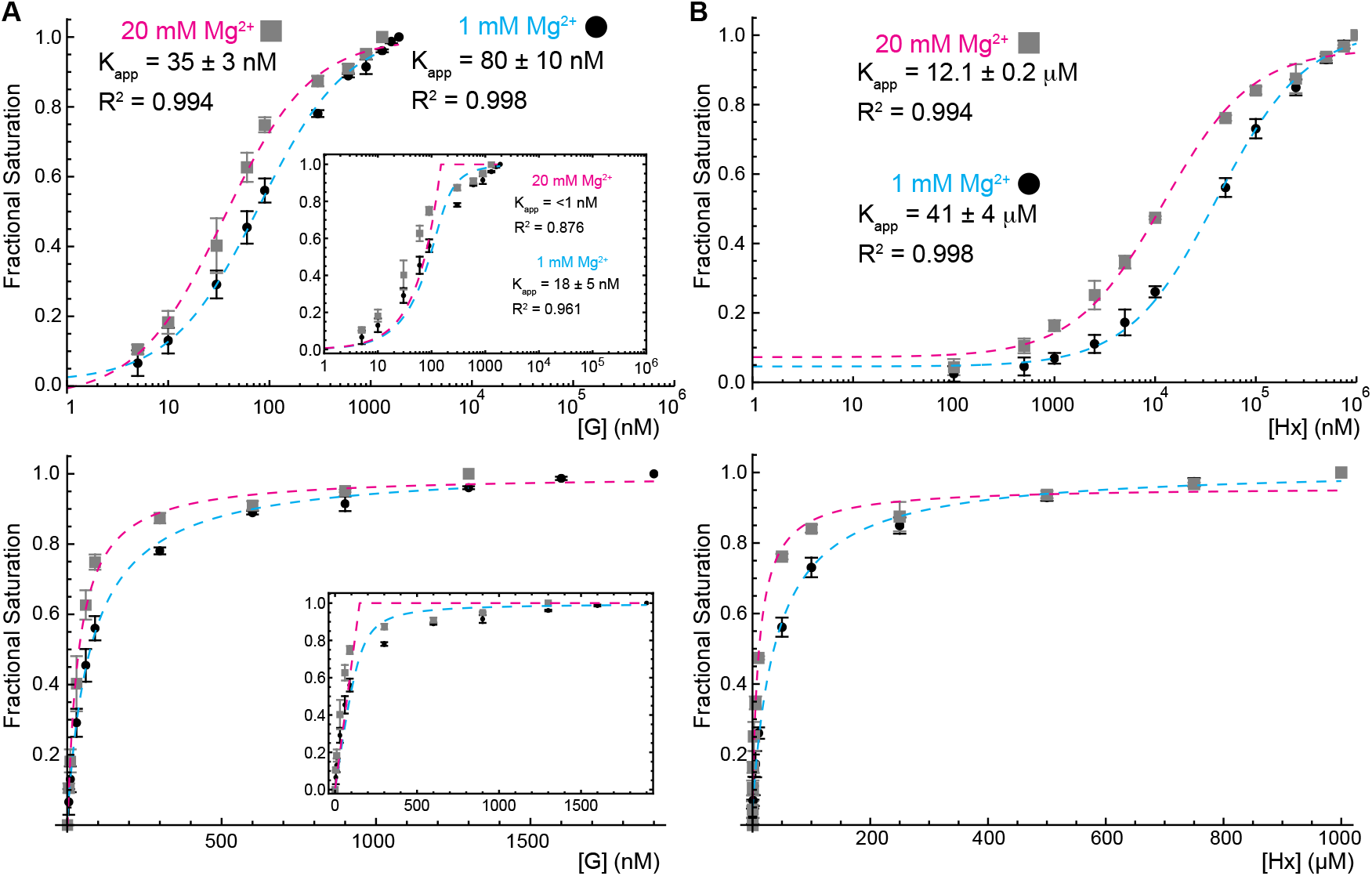
Ligand titrations. Error bars represent the standard deviation across 3 experimental replicates. (**A**) Titration of G into pC74 GR in phosphate buffer supplemented with 1 mM MgCl_2_ (black; fit in dashed blue) or 20 mM MgCl_2_ (gray; fit in dashed pink). Fractional saturation was computed from the peak intensity of the emission spectrum at each ligand concentration and then fitted with a 4-parameter logistic equation with the Hill coefficient fixed at 1 as a function of total ligand concentration. Top: logarithmic scale on x-axis. Bottom: linear scale on x-axis. Insets: Titration curves fitted with a quadratic model to account for ligand depletion (see Methods 3.3.1). (**B**) 4PL fits as in panel A but with Hx titrated into pC74 GR.

Indeed, analysis using a quadratic model that accounts for mass balance [20,64] suggests smaller K_D_ values under both physiological and high Mg^2+^ concentrations, though the much higher RNA concentration limits the precision of these estimates. Additionally, controls were carried out in which G or Hx was titrated into pC riboside (pCr). pCr exhibited no change in emission intensity under all conditions (Fig. S6), confirming that the decrease in fluorescence observed with pC74 is due to ligand binding rather than photodamage or nonspecific interactions between pC and ligand. In summary, while pC74 GR shows a modest reduction in ligand-binding affinity compared to unlabeled GR, it maintains high specificity, including under physiological conditions where the low Mg^2+^ concentration might exacerbate any defects induced by pC labeling. This preservation of key functions exhibited by unmodified GR suggested that pC74 GR could provide meaningful information about structure and ligand binding through further spectroscopic studies.

### 2.5. Conformational ensemble of pC74

TCSPC allows for a greater examination of the conformations that pC74 samples within the binding pocket and how they give rise to the distinct emission behaviors of the G- and Hx-bound complexes. All pC74 GR decays were well fit by a 3E model (χ^2^ = 1.17-1.23; details in Appendix A), returning short, intermediate, and long decay components with shifting lifetimes and population weights between conditions (Fig. 4C, D; Table 4). All conditions measured (RNA alone, +1 mM Mg^2+^, +4 µM G or +310 µM Hx) exhibit a 14 ns component, which is attributed to a base-stacked and unpaired state. In the absence of ligand, this stacked state is the predominant conformation (RNA alone, w=0.44; RNA+1 mM Mg^2+^, w=0.51). When saturating concentrations of ligand are introduced, this population decreases by an amount commensurate with the degree of hydrogen bonding to the ligand (G, w=0.21; Hx, w=0.36), as was observed in the model 21-mers when St1ApCA is hybridized to complementary strands with different bases opposite pC (Table 3). In addition, all conditions measured exhibit an intermediate length decay component, which changes in length from 7 ns in the ligand-free state to 5.5 ns in the presence of either ligand without meaningfully changing in population (w≈0.30). This is assigned as an unpaired and unstacked state, observed in the corresponding 21-mers at 5.5 ns. Both ligands induce a shift in population to a short decay component, attributed to the base-paired and base-stacked state. In the presence of G, this 2 ns component becomes the predominant population (w=0.45). In the presence of Hx, this population is equal in weight to the base-stacked and unpaired state (w=0.34) and, at 1.5 ns, is more rapidly decaying than that of the G-bound pC74 GR. Thus, the lifetime of this state is inversely related to the degree of pC’s hydrogen bonding, a pattern that was also observed in the model double-stranded 21-mers (Table 3).

**Table 4.**
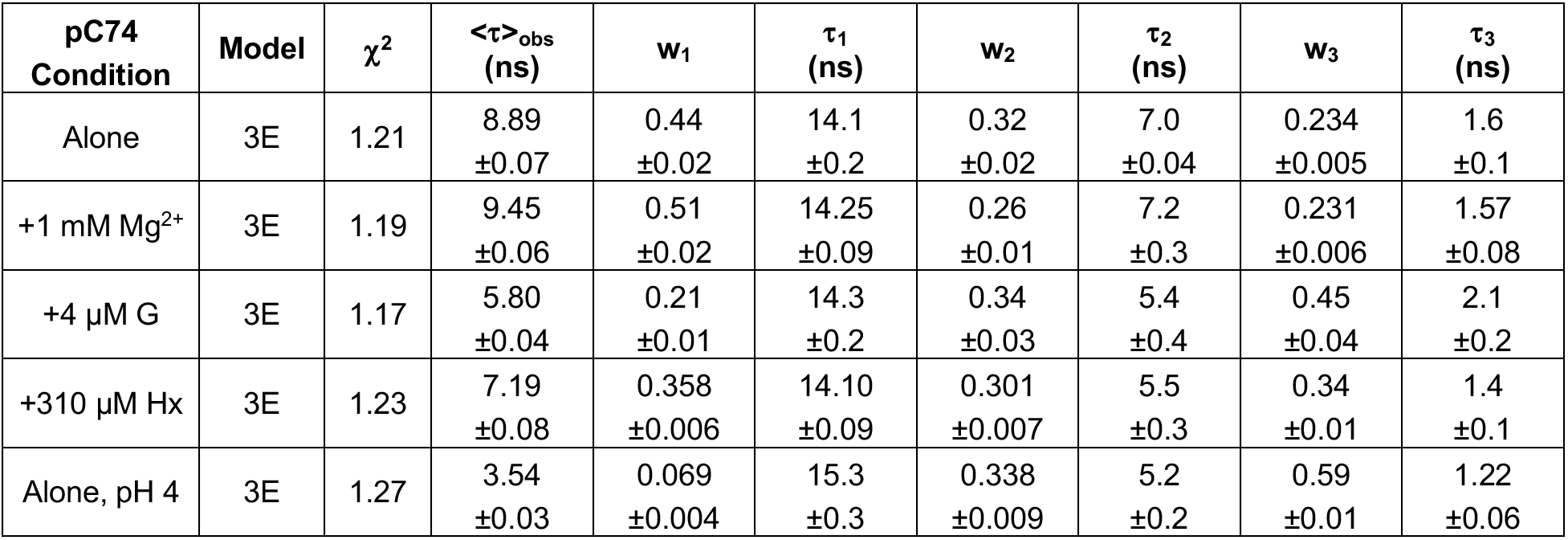
TCSPC fitting results for pC74 GR. Uncertainties represent the standard deviation across 3 successive data acquisitions. Unless otherwise stated, all data was collected at pH 7.4.

| pC74 Condition | Model | $\chi^2$ | $\langle\tau\rangle_{\text{obs}}$ (ns) | $w_1$ | $\tau_1$ (ns) | $w_2$ | $\tau_2$ (ns) | $w_3$ | $\tau_3$ (ns) |
| --- | --- | --- | --- | --- | --- | --- | --- | --- | --- |
| Alone | 3E | 1.21 | 8.89<br>$\pm 0.07$ | 0.44<br>$\pm 0.02$ | 14.1<br>$\pm 0.2$ | 0.32<br>$\pm 0.02$ | 7.0<br>$\pm 0.04$ | 0.234<br>$\pm 0.005$ | 1.6<br>$\pm 0.1$ |
| +1 mM Mg<sup>2+</sup> | 3E | 1.19 | 9.45<br>$\pm 0.06$ | 0.51<br>$\pm 0.02$ | 14.25<br>$\pm 0.09$ | 0.26<br>$\pm 0.01$ | 7.2<br>$\pm 0.3$ | 0.231<br>$\pm 0.006$ | 1.57<br>$\pm 0.08$ |
| +4 μM G | 3E | 1.17 | 5.80<br>$\pm 0.04$ | 0.21<br>$\pm 0.01$ | 14.3<br>$\pm 0.2$ | 0.34<br>$\pm 0.03$ | 5.4<br>$\pm 0.4$ | 0.45<br>$\pm 0.04$ | 2.1<br>$\pm 0.2$ |
| +310 μM Hx | 3E | 1.23 | 7.19<br>$\pm 0.08$ | 0.358<br>$\pm 0.006$ | 14.10<br>$\pm 0.09$ | 0.301<br>$\pm 0.007$ | 5.5<br>$\pm 0.3$ | 0.34<br>$\pm 0.01$ | 1.4<br>$\pm 0.1$ |
| Alone, pH 4 | 3E | 1.27 | 3.54<br>$\pm 0.03$ | 0.069<br>$\pm 0.004$ | 15.3<br>$\pm 0.3$ | 0.338<br>$\pm 0.009$ | 5.2<br>$\pm 0.2$ | 0.59<br>$\pm 0.01$ | 1.22<br>$\pm 0.06$ |

Unexpectedly, pC74 GR also exhibited a 1.5 ns decay component (w=0.24) in the absence of ligand. Though this timescale has typically been observed in base-paired contexts [31,34], it was also present in the mismatched dsRNA 1:2UUU (w=0.35, ***τ***=1.99; Table 3), demonstrating that a component of this length can arise without base pairing. In order to establish the conformation underlying this rapidly decaying population in pC74 GR, we performed measurements under acidic conditions to compare the fluorescence properties of denatured pC-labeled GR to those observed at pH 7.4 (Fig. 6). HCl has been used previously to rapidly denature RNA [65], and the concentration required to achieve denaturation of the GR (determined by CD to occur at pH 4; Fig. S7) altered the buffer conditions only by increasing the total chloride concentration from 100 mM to 117 mM. While the fluorescence decay of pC74 GR at pH 4 is still well fit by a 3E model (χ^2^=1.27), stark changes in the occupancy of the states described above are observed (Table 4). The 15 ns base-stacked population decreases drastically at pH 4 to a weight of only 0.07, while the shortest decay component becomes dominant (w=0.59) with a slight reduction in length to 1.1 ns (Fig. 6D). The increased population of this rapidly decaying state is reflected in a 67% decrease in the emission intensity (Fig. 6A). In contrast, pC71 GR exhibits a 67% increase in emission intensity at pH 4, resulting in a nearly identical emission intensity and lineshape to pC74 (Fig. 6B, C). This confirms that the strong emission of pC74, the weak emission of pC71, and its distinctive ∼530 nm shoulder all reflect unique environments within the native riboswitch structure. At pH 4, pC71 GR required a 3E fit (χ^2^=1.28), in contrast to the 1E1G_S_ model that was optimal under neutral conditions. As in pC74 GR, the shortest decay component (0.97 ns) dominates with a population weight of 0.57 (Table 2). This same shift is observed in St1ApCA, which also required a 3E fit under acidic conditions, as opposed to the 2E fit that was adequate under neutral conditions. Notably, the new component identified by the 3E model (1.5 ns; w=0.56) closely mimics both the lifetime and the population weight of the short component observed in pC71 GR and pC74 GR at pH 4.

**Figure 6.**
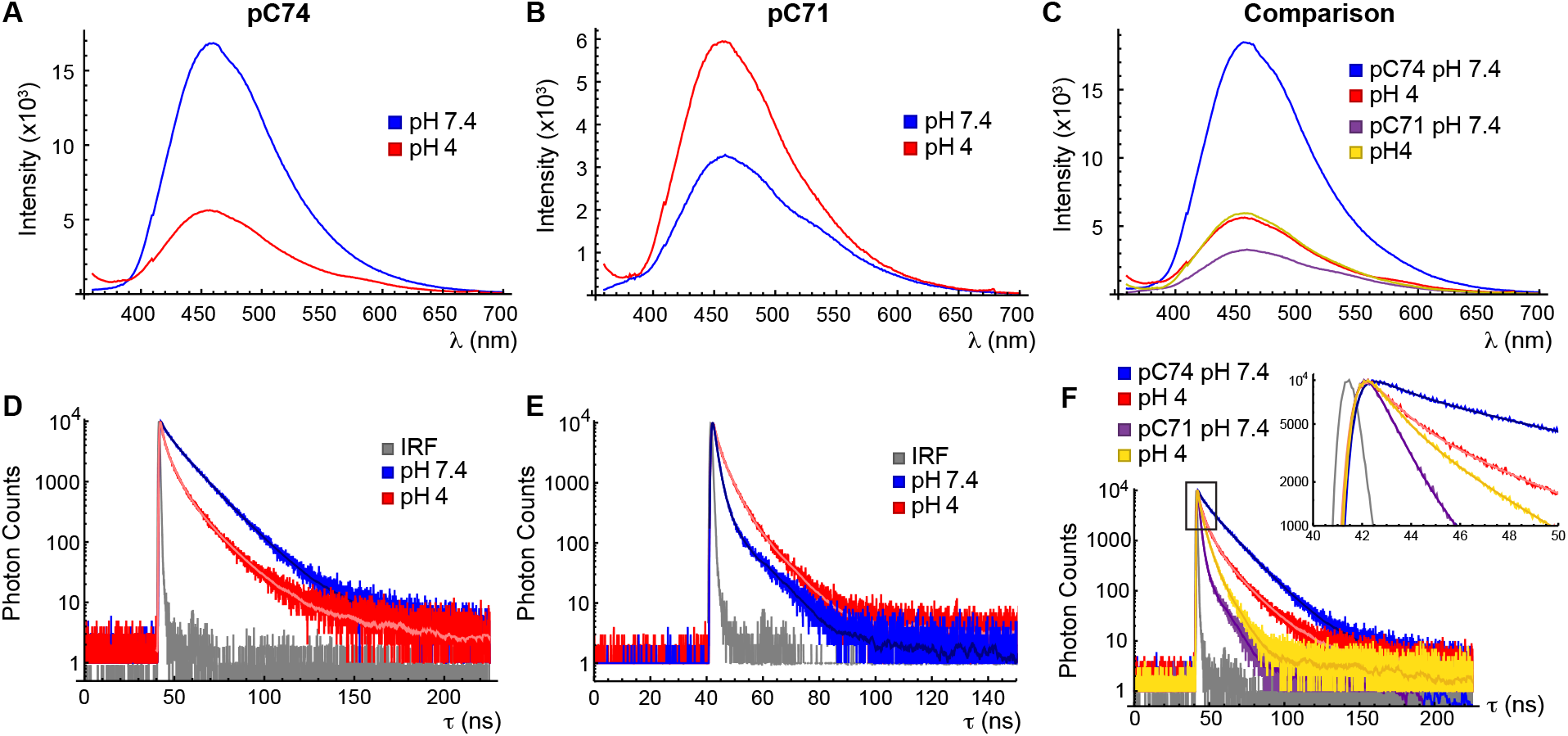
Steady-state and time-resolved spectroscopy of pC74 and pC71 GR under native (pH 7.4) and denaturing (pH 4) conditions. (**A**) Fluorescence emission spectra of pC74 in standard phosphate buffer at pH 7.4 (blue) or phosphate buffer adjusted to pH 4 using HCl (red). (**B**) Same as panel A for pC71 GR. (**C**) Overlaid emission spectra of pC74 GR at pH 7.4 (blue) and pH 4 (red) and pC71 GR at pH 7.4 (purple) and pH 4 (yellow). (**D**) Fluorescence decays of pC74 GR at pH 7.4 (blue) and pH 4 (red). 3E fits are overlaid in contrasting color. (**E**) Fluorescence decays of pC71 GR at pH 7.4 (blue) and pH 4 (red). 1E1G_S_ (pH 7.4) and 3E (pH 4) fits are overlaid in contrasting color. (**F**) Overlaid fluorescence decays of pC74 GR at pH 7.4 (blue) and pH 4 (red) and pC71 GR at pH 7.4 (purple) and pH 4 (yellow). A close-up of the short-time region is shown as an inset.

Free pCr exhibits a vastly dominant (w=0.99) lifetime of 1.8 ns under both neutral and acidic conditions (Table 3, Fig. S8), representing the characteristic lifetime of pC in a completely unstacked and unpaired environment. We therefore propose that under denatured conditions, pC71, pC74 and the pC label within St1ApCA are all able to significantly populate a “riboside-like” state of pC, more broadly termed an “extrahelical conformation” [66,67]. Extrahelical conformations have been observed for 2-AP in mis-matched [66] and fully complementary [68] DNA duplexes, as well as riboswitches [20]. As we propose here for pC, extrahelical conformations of 2-AP are distinguished by fluorescence lifetimes comparable to free nucleosides. As the lifetimes of pCr and the short component of pC74 GR are nearly independent of pH (Fig. S8; Table 3; Table 4), this suggests that in the absence of ligand under native conditions, pC74 is able to sample an extrahelical conformation in which it is completely flipped out from the J3/1 linker. This state is then overcome by the characteristic, higher population weight conformation of pC74 hydrogen-bonded to ligand. 1:2UUU is similarly able to sample an extrahelical conformation (the previously mentioned 2 ns decay component) that is overcome by the rapidly decaying base-paired states present in the fully complementary dsRNAs 1:2UGU and 1:2UIU. The presence of an extrahelical conformation of 1:2UUU, in addition to its substantial population of the 5.5 ns “unstacked” conformation, indicates that the mis-matched, AU-rich core of 1:2UUU has a locally weaker base-stacking stability than the other duplexes investigated here or previously [34]. As the lifetimes of the extrahelical and base-paired conformations of pC are similar to each other, control measurements and structural information obtained by complementary methods should be leveraged in order to accurately assign decay components with lifetimes of 0.5-2 ns to molecular conformations.

### 2.6. FDCD reveals ligand-induced quenching of base-stacked conformations

We next utilized CD and fluorescence-detected CD (FDCD) spectroscopy to investigate the local and global conformations adopted by pC74 GR in response to Mg^2+^ and ligand. The standard CD spectra revealed a lineshape characteristic of A-form dsRNA [69–71], with a prominent positive feature at ∼265 nm and a prominent negative feature at ∼210 nm, and no changes were observed upon addition of Mg^2+^, G or Hx (Fig. 7). This is consistent with the pre–organization of the riboswitch into a global structure dominated by three base-paired helical domains (P1, P2 and P3) [52]. While pC has the potential to generate a CD signal in its longest-wavelength band [71], this was unresolvable in the standard CD spectra due to its low amplitude and partial spectral overlap with the strong CD signal generated by the 68 native bases in pC74 GR (Fig. 7). CD was therefore unable to report on the local environment around the labeling site.

**Figure 7.**
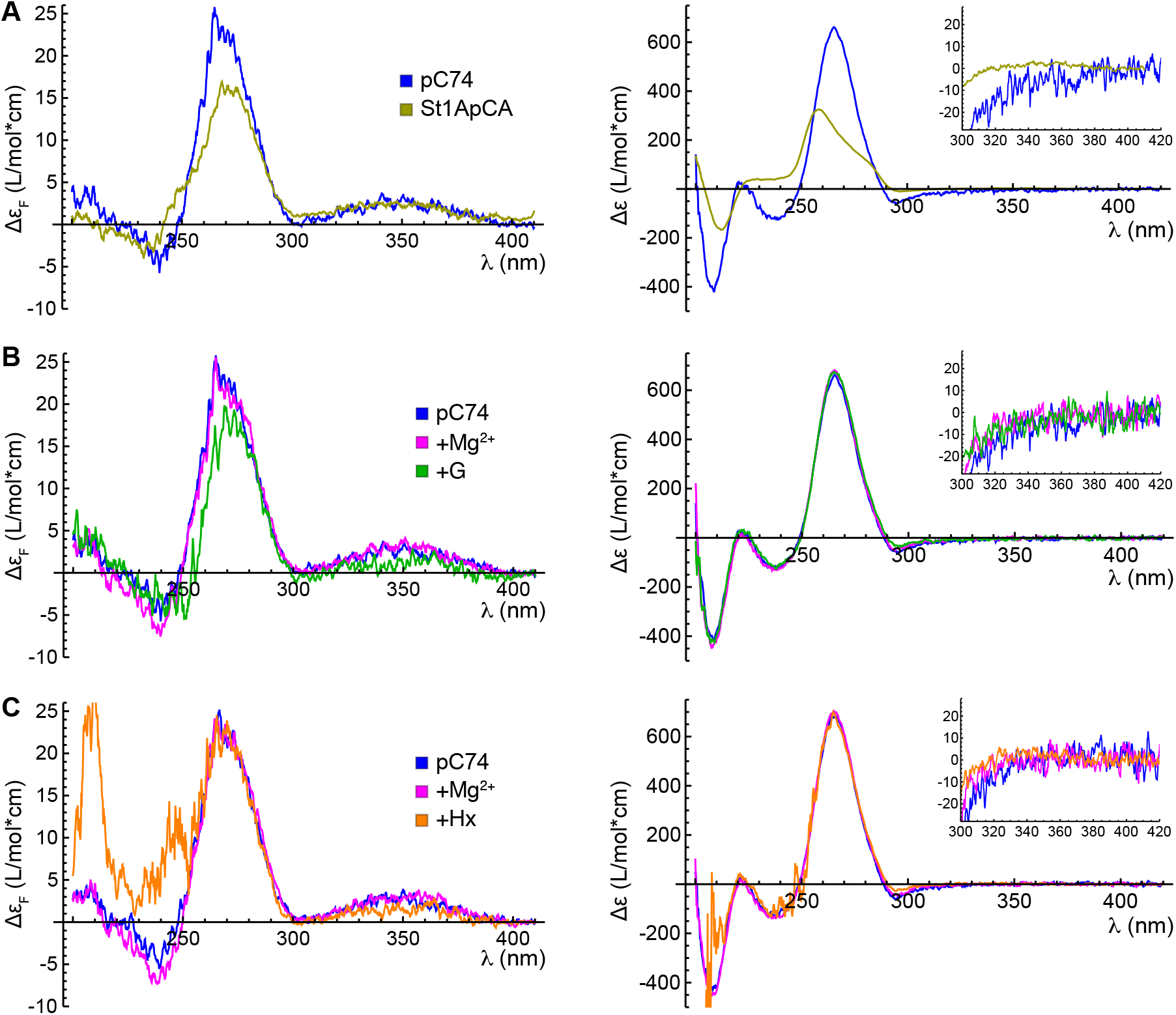
CD and FDCD spectroscopy of pC74 GR. (**A**) FDCD (left) and CD (right) spectra of pC74 GR (blue) and St1ApCA (yellow) in standard phosphate buffer. The inset shows a close-up of the long-wavelength region. (**B**) FDCD (left) and CD (right) spectra of pC74 GR in standard phosphate buffer (blue), buffer supplemented with 1 mM MgCl_2_ (magenta) or buffer supplemented with 1 mM MgCl_2_ and 4 µM G (green). (**C**) FDCD (left) and CD (right) spectra of pC74 GR in standard phosphate buffer (blue), buffer supplemented with 1 mM MgCl_2_ (magenta) or buffer supplemented with 1 mM MgCl_2_ and 310 µM Hx (orange).

FDCD reveals the CD signals that arise from conformations of an RNA that exhibit bright pC fluorescence. These signals can be generated by electronic transitions on pC itself, transitions delocalized across pC and native bases, and energy transfer from native bases to pC [73]. As previously observed in pC-labeled trinucleotides and ssRNA [34], the FDCD spectra of St1ApCA and pC74 GR both exhibit a positive feature in the longest-wavelength absorption band of pC and a sigmoidal feature at shorter wavelength arising from excitation of nearby native bases followed by energy transfer to pC. The spectra differ in their details, however, due to the unique structure of the binding pocket surrounding pC74. Most notably, the intensity of the short-wavelength feature is higher in pC74 GR than in St1ApCA, suggesting that a larger number of bases participate in energy transfer to pC74, and its lineshape also differs, reflecting the difference in structure surrounding pC (Fig. 7A).

The intensity of a short-wavelength negative feature becomes slightly but reproducibly larger upon addition of Mg^2+^ to pC74 GR, and the intensity of the entire FDCD spectrum becomes significantly lower upon addition of a saturating concentration of G (Fig. 7B). This shows that binding of G has the effect of quenching emission from the conformational subpopulations in which pC and nearby native bases have the strongest CD signals, while subpopulations with weaker CD remain bright. TCSPC revealed that a dominant effect of ligand is to shift population from a base-stacked but unpaired conformation to a stacked and paired conformation (Table 4). Thus, the ensemble of base-stacked conformations becomes dominated by the darker, ligand-bound state in the presence of G, consistent with the reduction in FDCD intensity upon ligand binding. Addition of Hx induces a smaller reduction in FDCD than G in the longest wavelength band, consistent with the smaller shift of population from bright to dark base-stacked conformations observed by TCSPC. Changes also occur in the short-wavelength feature, but their nuances are likely distorted by imperfect correction for the strong inner filter effect generated by saturating concentrations of Hx (Fig. S4), which also impacted the fluorescence excitation spectra discussed above.

### 2.7. Stopped-flow fluorescence spectroscopy reveals differences in ligand association rates

Uncertainty of k_assoc_ reported (±0.2*10^4^ M^-1^s^-1^) is the standard error of the slope obtained from the linear fit. (**C**) Corrected fluorescence traces for the association of 50 µM (purple), 125 µM (blue), 250 µM (yellow), or 500 µM (red) Hx to pC74 GR. (**D**) Observed rate constant plotted as a function of Hx concentration. A linear fit (red) returned an association rate constant of 0.03*10^4^ M^-1^s^-1^. Uncertainty of k_assoc_ reported (±0.003*10^4^ M^-1^s^-1^) is the standard error of the slope obtained from the linear fit.

Previous studies have utilized 2-AP as a ligand analogue for fluorescence studies of GR variants, exploiting the fact that their affinities for 2-AP were proportional to, albeit much weaker than, their affinities for G [42]. Furthermore, 2-AP has been used as a ligand analogue in stopped-flow fluorescence studies of ligand binding to a naturally occurring adenine-sensing riboswitch [40] and a mutant of GR designed to preferentially bind adenine (C74U) [41]. In order to determine if pC could be utilized for kinetic studies of ligand binding, we carried out stopped-flow fluorescence measurements on pC74 GR where increasing concentrations of either G or Hx were introduced (Fig. 8). The ligand-dependent quenching of pC74 allowed observed association rates to be determined from the decrease in fluorescence intensity over time. All measurements were carried out under pseudo-first order conditions, with pC74 GR at 55 nM and ligand in a large excess (275 to 2750 nM G; 50 to 500 µM Hx). The fluorescence traces were scaled to run from a maximum of 1 to a minimum of 0. These corrected traces were fitted with a bi-exponential function, and the observed association rates (k_obs_) were calculated as a weighted average of the fast and slow decay components’ rate constants (Fig. S9, individual weights and rate constants in Table S1). As previously done for 2-AP binding to C74U GR [41], the association rate (k_assoc_) was estimated as the slope of a linear fit to k_obs_ vs. ligand concentration (Fig. 8). pC74 exhibited clear time-dependent fluorescence quenching upon introduction of both G and Hx. Far slower changes were observed when buffer was introduced or when ligand was introduced to pCr (Fig. S9C-E), indicating that quenching of pC74 emission after rapid mixing arises from ligand binding. As expected, Hx required much higher concentrations to achieve the same degree of fluorescence quenching as G, which is reflected in their k_assoc_ values: G=4.1x10^4^ M^-1^s^-1^ (standard error from linear fit: 0.2x10^4^ M^-1^s^-1^); Hx = 0.033x10^4^ M^-1^s^-1^ (standard error: 0.003x10^4^ M^-1^s^-1^). Interestingly, the k_assoc_ value of G is nearly identical to the rate determined for 2-AP binding to C74U GR at 1.25 mM Mg^2+^ (4.4x10^4^ M^-1^s^-1^) [41]. The necessity of a bi-exponential fit may reflect a previously proposed binding model in which GR samples “closed” and ”open” states, where only the latter is capable of binding ligand [41,42]. These results confirm that pC substitutions can be used to reliably measure binding kinetics between RNA and small-molecule ligands.

**Figure 8.**
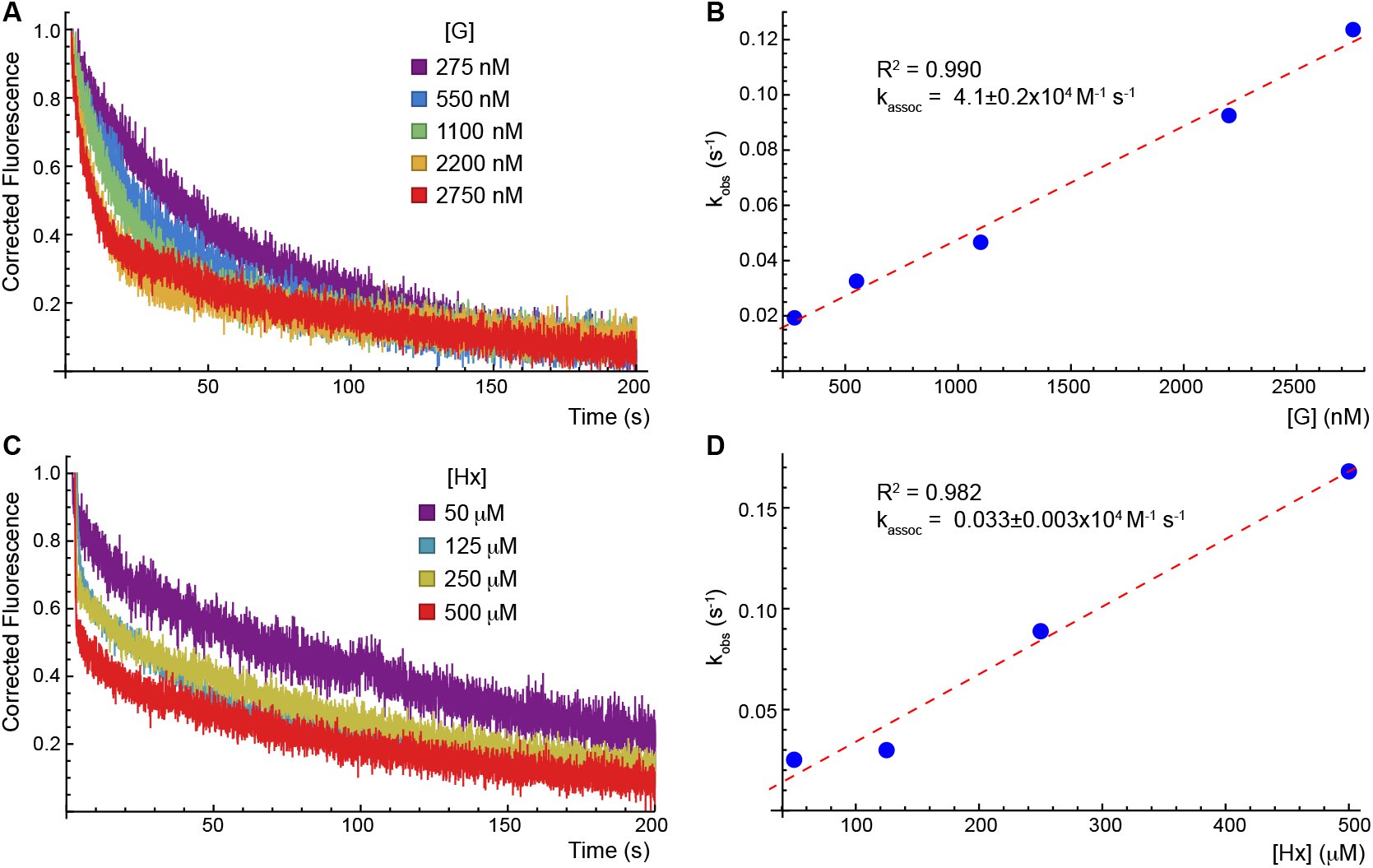
Stopped-flow fluorescence spectroscopy. (**A**) Corrected fluorescence traces for the association of 275 nM (purple), 550 nM (blue), 1100 nM (green), 2200 nM (yellow) or 2750 nM (red) G to pC74 GR. Raw fluorescence was corrected to run from a maximum of 1 (representing the fluorescence of ligand-free pC74 GR) to a minimum of 0 (fluorescence of ligand-bound pC74 GR). See Methods for data processing details. (**B**) Observed rate constant plotted as a function of G concentration. A linear fit (red) returned an association rate constant of 4.1*10^4^ M^-1^s^-1^.

## 3. MATERIALS AND METHODS

### 3.1. Reagents

RNA oligonucleotides were purchased from Horizon Discovery (Dharmacon) and DNA oligonucleotides were purchased from Integrated DNA Technologies (IDT) (Table 1) and deprotected and desalted by the manufacturer. The GR RNA sequence was split into a 5’ and a 3’ segment. pC71, pC74 and unmodified versions of the 3’ segment were all purchased with a 5’ phosphate. In addition, pC-labeled 3’ segments and all 21-mers were high-pressure liquid chromatography-purified by the manufacturer. Oligonucleotide stock solutions were prepared in DEPC-treated water and the A260 was used to determine the concentration using the extinction coefficient provided by the manufacturer. pCr was obtained from Biosearch Technologies (PYA11092-B010) and prepared in MilliQ H_2_O at a concentration of 5 mM. G was obtained from Sigma Aldrich (G11950-10G) and prepared in 0.4 M HCl at a concentration of 3 mM. Hx was obtained from Sigma Aldrich (H9377-5G) and prepared in 50 mM NaOH at a concentration of 53 mM. Buffers used for spectroscopy were prepared in MilliQ H_2_O and sterile-filtered prior to use.

### 3.2. Riboswitch Preparation

Full-length GR samples were prepared via splint-mediated ligation utilizing T4 RNA ligase 2 (Molecular Cloning Laboratories T4RL2-100). Five nanomoles of each of the 5’ segment, the 3’ segment, and DNA splint (Table 1) were combined and heated at 90 °C for two minutes and cooled at RT for 10 minutes. 10x T4 RNA ligase 2 buffer (500 mM Tris-HCl, 20 mM MgCl_2_, 10 mM DTT, pH 7.5 @ 25 °C) was to a concentration of 1x 30 seconds into cooling. Once cooled, sufficient buffer was added to reach a final concentration of 1x in the complete reaction, ATP to a final concentration of 200 µM and water to a final volume of 150-175 µL as required to accommodate the volumes of oligonucleotide. After mixing, 60 units of T4 RNA ligase 2 were added. The reactions were incubated at RT for 3 hours, then quenched with 2x formamide loading buffer (90% formamide, 1x TBE) and stored at -20 °C overnight. The reactions were then run on a 10% polyacrylamide, 8 M urea gel. The RNA was recovered from the product band via the crush and soak method or electroelution using a BioRad model 422 electro-eluter, followed by ethanol precipitation in the presence of 300 mM NaOAc.

### 3.3. Spectroscopy

Unless otherwise noted, all spectra were collected at 20 °C in a buffer consisting of 20 mM NaiPO4 (pH 7.4) and 100 mM NaCl (termed “standard buffer”). GR and 21-mers were annealed before measurement by heating at 90 °C for two minutes then cooled at RT for 10 minutes. Where noted, MgCl_2_, G and/or Hx were added after cooling. Experiments under acidic conditions were performed by preparing samples in standard buffer and then adding HCl to achieve a pH of 4. Samples were incubated at RT for five minutes after addition of ligand or other solvent components. Analysis was carried out in Mathematica (Wolfram Alpha, v12.3.1.0) unless otherwise stated.

#### 3.3.1. Steady-state absorbance and fluorescence spectroscopy

Steady-state spectra were collected on an Edinburgh FS5 spectrofluorometer with a 1 nm step size and a 1 s dwell time. Fluorescence titrations were collected with GR or pCr at a concentration of 150 nM. The fractional saturation (FS) was computed from the peak fluorescence intensity (I_max_) at each ligand concentration (Equation 1), plotted as a function of total ligand concentration, and fitted with a 4PL (Equation 2; Hill coefficient fixed at 1) or quadratic (Equation 3) binding model. Reproducibility suffered at RNA concentrations lower than 150 nM due to low emission intensity. As such, apparent K_D_ values below or around 150 nM obtained from a 4PL model represent the maximum possible value.

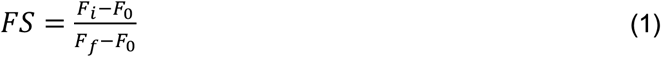

In equation 1, *F_i_* represents I_max_ value at ligand concentration *i*, F_0_ represents the I_max_ value in the absence of ligand, and *F_f_* represents the I_max_ value at the highest measured ligand concentration.

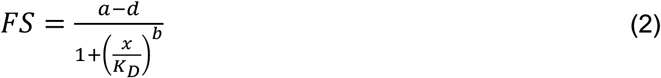

In equation 2, a and d represent the minimum and maximum values of the curve and b represents the Hill coefficient.

In equation 3, *R* represents the RNA concentration and *L* represents the ligand concentration.

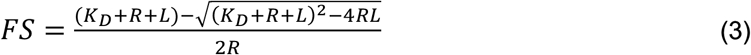

All other steady-state spectra were collected at an RNA concentration of 1 µM. Emission spectra were recorded by exciting at 340 nm with a 5 nm bandwidth while scanning from 360 to 700 nm with an emission bandwidth of 5 nm for fluorescence titrations or 2 nm for all other steady-state spectra.

Excitation spectra were recorded by monitoring emission at 460 nm with a 5 nm bandwidth while scanning from 220 to 420 nm with a 2 nm excitation bandwidth. 21-mer and GR excitation spectra were corrected for the inner filter effect using absorbance spectra collected under the same conditions.

#### 3.3.2. TCSPC

TCSPC measurements were collected on a Horiba Fluoromax-5 with a TemPro fluorescence lifetime system equipped with a 340 nm NanoLED excitation source (1 MHz repetition rate). Fluorescence decays were collected to a peak count of 10,000 across a 200 ns window with bins of 55 ps. A 420 nm long-pass filter was placed in the emission path while neutral density filters were placed in the excitation path with sufficient optical density to avoid pulse pile up. Instrument response functions (IRFs) were collected by acquiring data on buffer with no long-pass filter in place.

Decays were analyzed by iterative reconvolution fitting in Matlab (R2024a) using open-source software [73], as previously described [34]. Upon fitting with a constant IRF background of 0 counts, a timing shift between the IRF and decays of ∼50 ps was returned. Under ideal conditions, decays with time constants of ∼1/10 the full width at half maximum (FWHM) of the IRF can be recovered, with the lower limit determined by the bin width and timing shift. As the FWHM of the IRF was ∼850 ps in the measurements reported here, the shortest components (∼600 ps) are well within this lower limit. The observed average lifetime was calculated as a population weighted average of the lifetimes obtained from exponential or gamma distribution parameters (Equation 4).

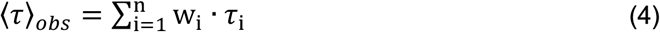

A gamma component’s time constant was defined as the mean of the gamma distribution (Equation 5), as in analysis of other FBAs [62].

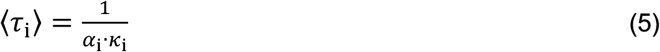

#### 3.3.3. CD and FDCD

CD and FDCD spectra were recorded as previously described [34,74] on a Jasco J1500 circular dichroism spectrometer with an FDCD-551 attachment and a 420 nm long-pass filter placed in front of the emission detector. FDCD scans were collected at an excitation bandwidth of 4 nm, a scanning speed of 20 nm/min with 4 s digital integration time over the range 200 – 410 nm. The gain on the emission detector was set such that the maximum signal in the DC measurement channel was ∼1 V. FDCD spectra of GR were collected with RNA at a concentration of 200 nM, requiring 20 accumulations to be collected and averaged for sufficient signal-to-noise ratio. St1ApCA was measured at a concentration of 1 µM, requiring 10 accumulations. CD scans were collected under the same conditions out to 420 nm, only requiring 3 accumulations.

CD spectra were background-corrected using buffer scans and converted to units of Δε (L/mol*cm). FDCD spectra were background-corrected using sodium fluorescein in water at concentrations allowing for similar detector gain to each pC sample [34,76]. Sample and fluorescein scans were corrected with buffer and water, respectively. *_θ_*_F_, was calculated by dividing the FDCD channel by the DC channel. *_θ_*_F_ of fluorescein was smoothed with a Weiner filter (r=15) and then subtracted from *_θ_*_F_ of the corresponding sample.

Established theory [73] was used to convert *_θ_*_F_ to the quantity Δε_F_ (Equation 6), analogous to the typical and readily interpretable units of CD, Δε (L/mol*cm).

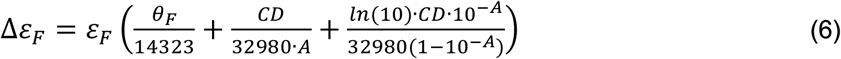

ε_F_ was approximated by dividing the DC signal at each wavelength by the measured Xe lamp intensity and then corrected for the inner filter effect using the corresponding absorbance spectrum. These excitation spectra were converted to units of L/mol*cm by scaling the value at 347 nm to match that as previously reported [77].

#### 3.3.4. Stopped-flow fluorescence spectroscopy

Stopped-flow fluorescence measurements were performed on an Edinburgh FS5 spectrofluorometer using an SFA-20 Rapid Kinetics Accessory equipped with a pneumatic drive attachment. RNA was prepared at a concentration of 110 nM in standard buffer + 1 mM MgCl_2_ resulting in a final concentration of 55 nM after mixing. Ligands were prepared at 2x the desired final concentration in standard buffer + 1 mM MgCl_2_. Fluorescence intensity was monitored at 460 nm with a 10 nm bandwidth under continuous excitation at 340 nm with a 10 nm bandwidth. Traces were collected over 200 s (G) and 300 s (Hx) with a time resolution of 30 and 40 ms, respectively. Each line of the flow circuit was rinsed with 10 mL of water before priming with its corresponding RNA, buffer, or ligand solution. The ligand line was rinsed with 10 mL of water in between ligand concentrations.

Fluorescence intensity traces were corrected to run from 1.0 to 0 (Equation 7) before fitting with a bi-exponential function (Equation 8). Quality of fit was assessed by R^2^ and the results are reported in Table S1. The observed rate constant (k_obs_) was calculated as a weighted average of the two rate constants (Equation 9). *A*_i_ represents the raw amplitude returned from the bi-exponential fit, *k*_i_ represents the corresponding rate component, and *I_0_* represents a constant y-offset.

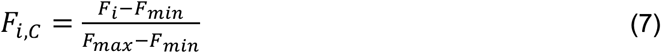

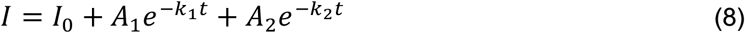

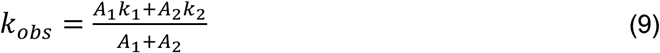

### 3.4. Computational methods

All calculations were carried out in Gaussian09 [78]. Methods and basis sets were selected based on the recommendations for pC-containing systems presented in Nguyen et al. [79]. First, the ground state molecular geometries of A, U, and pC nucleobases were optimized at the MP2/6–31G(d,p) level.

Using previously published coordinates for A-form trinucleotides [32] or the corresponding GR crystallographic coordinates (PDB: 4FE5 (ApCA) [45], 2EES (ApCU) [53]), the optimized monomers were assembled into ApCA, ApCU, and UpCU trimers for further calculations. Configuration interaction singles with perturbative second order corrections [CIS(D)] at the aug-cc-pVDZ level of theory was used to calculate the vertical excitation energy of the first 10 excited states. For each excited state, the natural transition orbitals were calculated at the same level of theory.

## 4. CONCLUSIONS

In summary, we have demonstrated the use of pC as a multifaceted, minimally perturbing probe that can report on the conformations sampled by specific nucleotides within a riboswitch and can act as a direct sensor of ligand binding. The ligand-dependent quenching of pC74 enabled the analysis of many aspects of GR structure and function, such as its affinity for various ligands, the conformational ensemble within the binding pocket, and the kinetics of ligand binding. We show that pC substitution preserves essential properties of the GR even when it replaces a critical ligand-binding residue, such as distinct binding affinity and kinetics determined by the number of hydrogen bonds formed with the ligand. We show that pC can sample base-stacked, unstacked, extrahelical and hydrogen-bonded conformations within the complex environment of the binding pocket, which exhibit varying degrees of fluorescence quenching or enhancement. These findings establish a framework for the broader use of pC in complex RNAs of biological and technological interest.

## Supplementary Materials

Figure S1: Natural transition orbitals of S0-S1 transition.; Figure S2: Normalized fluorescence emission and excitation spectra.; Figure S3: CD spectra of pC-labeled and unlabeled GR.; Figure S4: Absorbance spectra of G and Hx in a 1 cm pathlength cuvette at the concentrations used in this study.; Figure S5: Emission spectra of pC74 GR used to construct titration curves in Fig. 5.; Figure S6: Control titrations showing minimal photobleaching or nonspecific response of pC to G or Hx.; Figure S7: CD spectra of unmodified GR at varying pH.; Figure S8: Acid denaturation controls.; Figure S9: Stopped-flow fluorescence spectroscopy fitting and controls.; Table S1: Bi-exponential fitting results for stopped-flow fluorescence spectroscopy.

## Author Contributions

Conceptualization, T.L.C. and J.R.W.; methodology, T.L.C.; formal analysis, T.L.C.; investigation, T.L.C. and G.V.; writing—original draft preparation, T.L.C. and J.R.W.; writing—review and editing, G.V.; visualization, T.L.C. and J.R.W.; supervision, J.R.W.; project administration, J.R.W.; funding acquisition, J.R.W. All authors have read and agreed to the published version of the manuscript.

## Data Availability Statement

The original contributions presented in this study are included in the article/supplementary material. Further inquiries can be directed to the corresponding author.

## Abbreviations

The following abbreviations are used in this manuscript:

FBAs: fluorescent base analogues
TCSPC: Time-correlated single-photon counting
pC: pyrrolocytosine
CD: circular dichroism
pCr: pC riboside
FDCD: fluorescence-detected CD
2-AP: 2-aminopurine
CT: charge transfer
GR: guanine riboswitch
QM: quantum mechanical
G: guanine
ss: single-stranded
Hx: hypoxanthine
ds: double-stranded
WCF: Watson-Crick-Franklin

## APPENDIX A: Selection of fitting models for TCSPC

For the complex RNAs reported in this work, we surveyed models containing 2-4 exponential components, as well as gamma (γ) components, which ascribe a decay component to a distribution of closely related structures [62]. We use the notation “xEyG_L_(_S_)” to concisely indicate a model containing “x” exponential decay components and “y” γ distribution components. L(S) indicates that the γ distribution’s mean is longer (shorter) than the exponential component’s time constant. An ideal fitting model will have a reduced χ^2^ value close to 1 and have unstructured residuals. χ^2^ values below 1.3 are generally considered to be acceptable, though values below 1.2 were obtainable for all pC-containing model oligonucleotides previously investigated by our lab [34]. For the data reported here, however, χ^2^ values of 1.2-1.3 were often observed across a range of models of varying complexity. For example, the single-stranded 21-mer St1ApCA exhibited an average χ^2^ of 1.27 when fitted with a 2E model, which improved only marginally to 1.25-1.26 across a wide range of more complex models (Table A1).

Because the χ^2^ values were marginal (1.2-1.3) under all tested models in certain cases, we considered the residuals more closely. An ideal model will have residuals that lack autocorrelation and are randomly distributed above and below zero. These properties can be quantified with the Durbin-Watson (DW) parameter and Runs Z test, respectively. Residuals are considered uncorrelated at the 95% confidence level if the DW parameter is between 1.75 and 2.25, with a value of 2 indicating no autocorrelation. Residuals are considered random at the 95% confidence level if the Z parameter is between -1.96 and 1.96, with the ideal value being 0 [74].

Below, we compare the χ^2^, Durbin-Watson and Z values across six models that were used to fit the decays of St1ApCA at pH 7.4. It is evident that models of complexity beyond 2E yield only small improvements in χ^2^, and that all models exhibit acceptable DW and Z values. 2E, 1E1G_S_, and 2G models were in agreement about the behavior of this RNA, all showing a short component of population weight ∼0.3 and lifetime ∼4-5 ns, and a long component of lifetime ∼12 ns accounting for the remainder of the population weight. We therefore selected the simple 2E model as representative of this family of behavior.

**Table A1.**
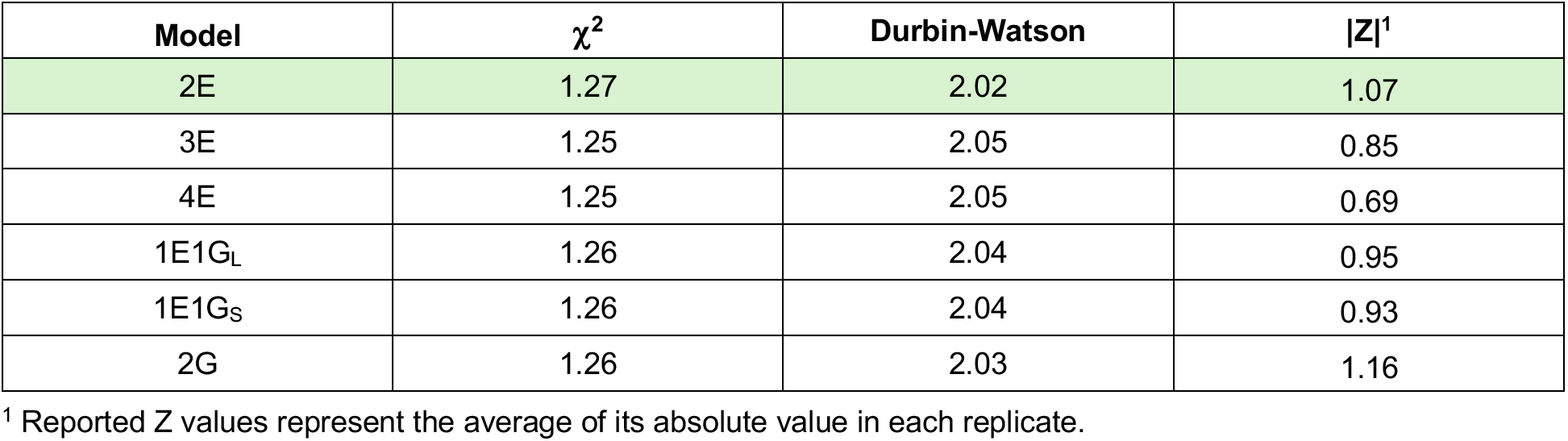
χ^2^, Durbin-Watson parameter and Runs test Z value for different models fit to decays of St1ApCA. Values represent the average across three successive data accumulations.

| Model | $\chi^2$ | Durbin-Watson | \|Z\|<sup>1</sup> |
| --- | --- | --- | --- |
| 2E | 1.27 | 2.02 | 1.07 |
| 3E | 1.25 | 2.05 | 0.85 |
| 4E | 1.25 | 2.05 | 0.69 |
| 1E1G<sub>L</sub> | 1.26 | 2.04 | 0.95 |
| 1E1G<sub>S</sub> | 1.26 | 2.04 | 0.93 |
| 2G | 1.26 | 2.03 | 1.16 |

<sup>1</sup> Reported Z values represent the average of its absolute value in each replicate.

For the three double-stranded 21-mers investigated here, 1:2UGU, 1:2UIU and 1:2UUU, the 2E fit was notably poor and all other models tested yielded a significant improvement in χ^2^. 3E was significantly better than 2E, 1E1G_S_ or 1E1G_L_ for 1:2UIU, was somewhat better for 1:2UUU, and was roughly comparable for 1:2UGU. For ease of comparison, we therefore selected 3E models for all double-stranded 21-mers. This contrasts with the previously investigated double-stranded 21-mers, which contained pC between two C residues opposite either GGG (fully complementary) or GUG (mis-matched) and were well fit by 2E models (χ^2^ = 1.18 and 1.13, respectively) [34].

**Table A2.**
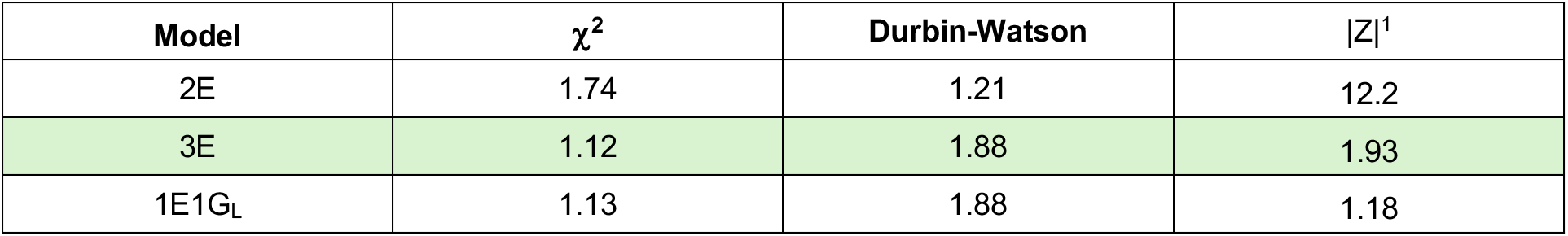
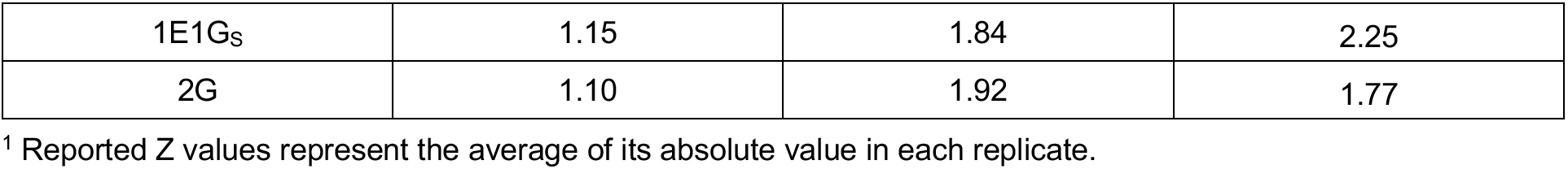
χ^2^, Durbin-Watson parameter and Runs test Z value for different models fit to decays of 1:2UGU. Values represent the average across three successive data accumulations.

**Table A3.** χ^2^, Durbin-Watson parameter and Runs test Z value for different models fit to decays of 1:2UIU. Values represent the average across three successive data accumulations.

| Model | $\chi^2$ | Durbin-Watson | \|Z\|<sup>1</sup> |
| --- | --- | --- | --- |
| 2E | 1.96 | 1.26 | 8.53 |
| 3E | 1.27 | 1.91 | 0.85 |
| 1E1G<sub>L</sub> | 1.37 | 1.79 | 0.89 |
| 1E1G<sub>S</sub> | 1.42 | 1.74 | 1.93 |
| 2G | 1.27 | 1.92 | 0.99 |

<sup>1</sup> Reported Z values represent the average of its absolute value in each replicate.

**Table A4.** χ^2^, Durbin-Watson parameter and Runs test Z value for different models fit to decays of 1:2UUU. Values represent the average across three successive data accumulations.

| Model | $\chi^2$ | Durbin-Watson | \|Z\|<sup>1</sup> |
| --- | --- | --- | --- |
| 2E | 1.38 | 1.63 | 5.66 |
| 3E | 1.17 | 1.91 | 1.29 |
| 1E1G<sub>L</sub> | 1.22 | 1.85 | 2.18 |
| 1E1G<sub>S</sub> | 1.20 | 1.88 | 1.89 |
| 2G | 1.19 | 1.89 | 1.82 |

<sup>1</sup> Reported Z values represent the average of its absolute value in each replicate.

2E models were universally poor for pC71 and pC74 GR. For pC71 at pH 7.4, we selected a 1E1G_S_ model, which yielded the best ꭓ2 value and residuals characteristics under the four measurement conditions tested (alone, +Mg^2+^, +G and +Hx) (values for RNA alone shown in Table A5 as an example).

**Table A5.** χ^2^, Durbin-Watson parameter and Runs test Z value for different models fit to decays of pC71. Values represent the average across three successive data accumulations.

| Model | $\chi^2$ | Durbin-Watson | \|Z\|<sup>1</sup> |
| --- | --- | --- | --- |
| 2E | 1.48 | 1.40 | 6.90 |
| 3E | 1.11 | 1.87 | 1.55 |
| 1E1G<sub>L</sub> | 1.16 | 1.83 | 1.49 |
| 1E1G<sub>S</sub> | 1.10 | 1.90 | 0.90 |

<sup>1</sup> Reported Z values represent the average of its absolute value in each replicate.

pC74 also exhibited a significant improvement in χ^2^ upon transitioning from a 2E model to a 3E or distribution model. Under 3E, 1E1G_L_ and 1E1G_S_ models, χ^2^ remained above 1.2 but the DW and Z parameters both suggested that residuals were random (values for RNA alone shown in Table A6 as an example). A 3E model was selected among these as it exhibited a slightly better χ^2^ and residuals characteristics across the four con-ditions tested (alone, +Mg^2+^, +G and +Hx). Only one replicate of pC74 yielded physically reasonable parameter values under a 4E model, and the χ^2^ value for that replicate improved only to 1.21 from 1.22 for a 3E fit.

**Table A6.** χ^2^, Durbin-Watson parameter and Runs test Z value for different models fit to decays of pC74. Values represent the average across three successive data accumulations.

| Model | $\chi^2$ | Durbin-Watson | \|Z\|<sup>1</sup> |
| --- | --- | --- | --- |
| 2E | 1.35 | 1.82 | 2.17 |
| 3E | 1.21 | 2.02 | 0.52 |
| 1E1G<sub>L</sub> | 1.23 | 1.98 | 0.80 |
| 1E1G<sub>S</sub> | 1.23 | 1.98 | 0.59 |

<sup>1</sup> Reported Z values represent the average of its absolute value in each replicate.

