## Supplemental Information for "Pyrrolocytosine as a Fluorescent Sensor of RNA-Small Molecule Interactions"

Includes:

Supplementary Figures S1-S9

Supplementary Table 1

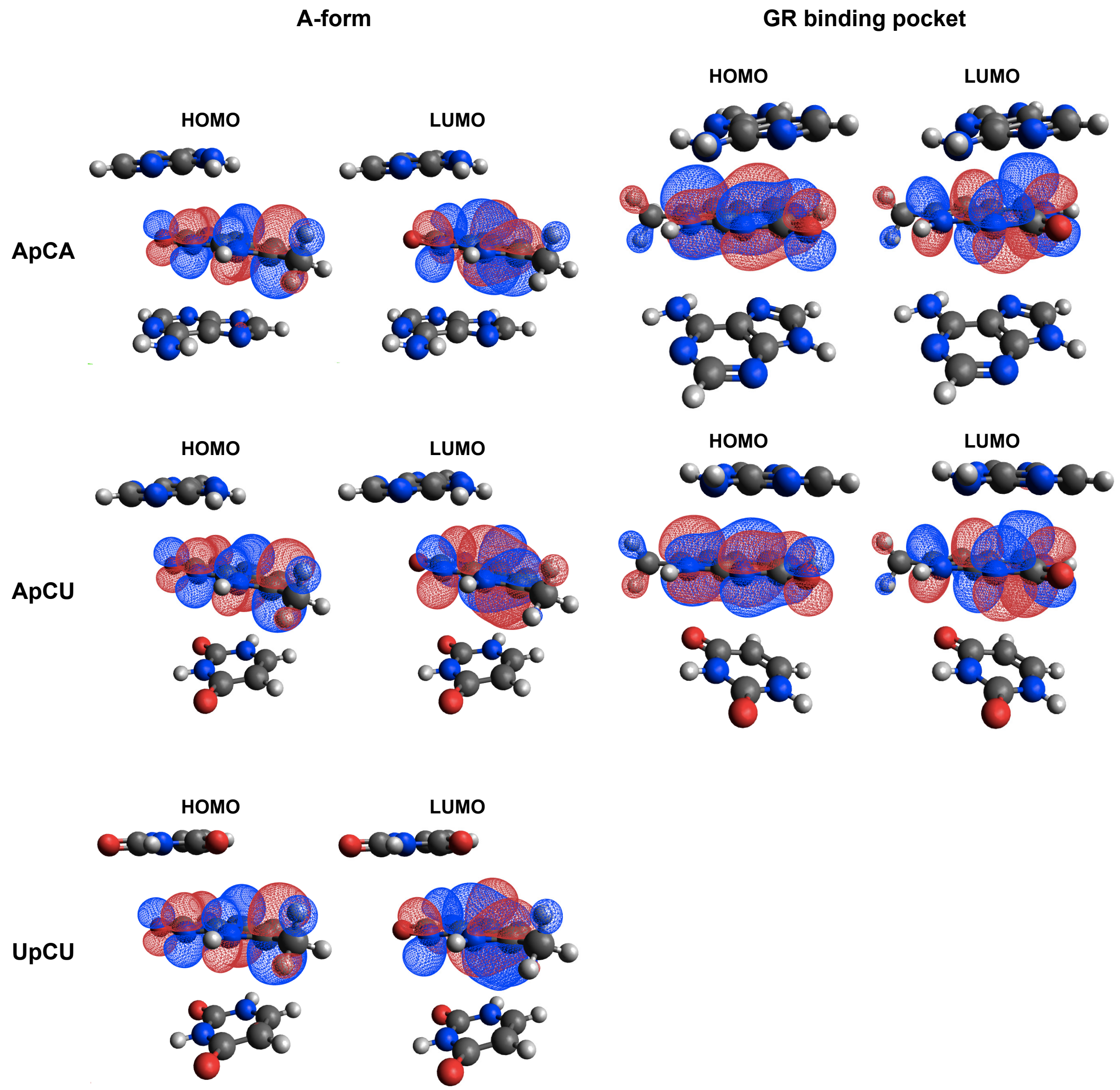


**Figure S1.** Natural transition orbitals of S0-S1 transition. Left column: ApCA (top), ApCU (middle), and UpCU (bottom) trimers in an A-form helical geometry. Right column: ApCA (top) and ApCU (bottom) trimers in the geometry of the GR binding pocket (ApCA PDB ID 2EES; ApCU PDB ID 4FE5).

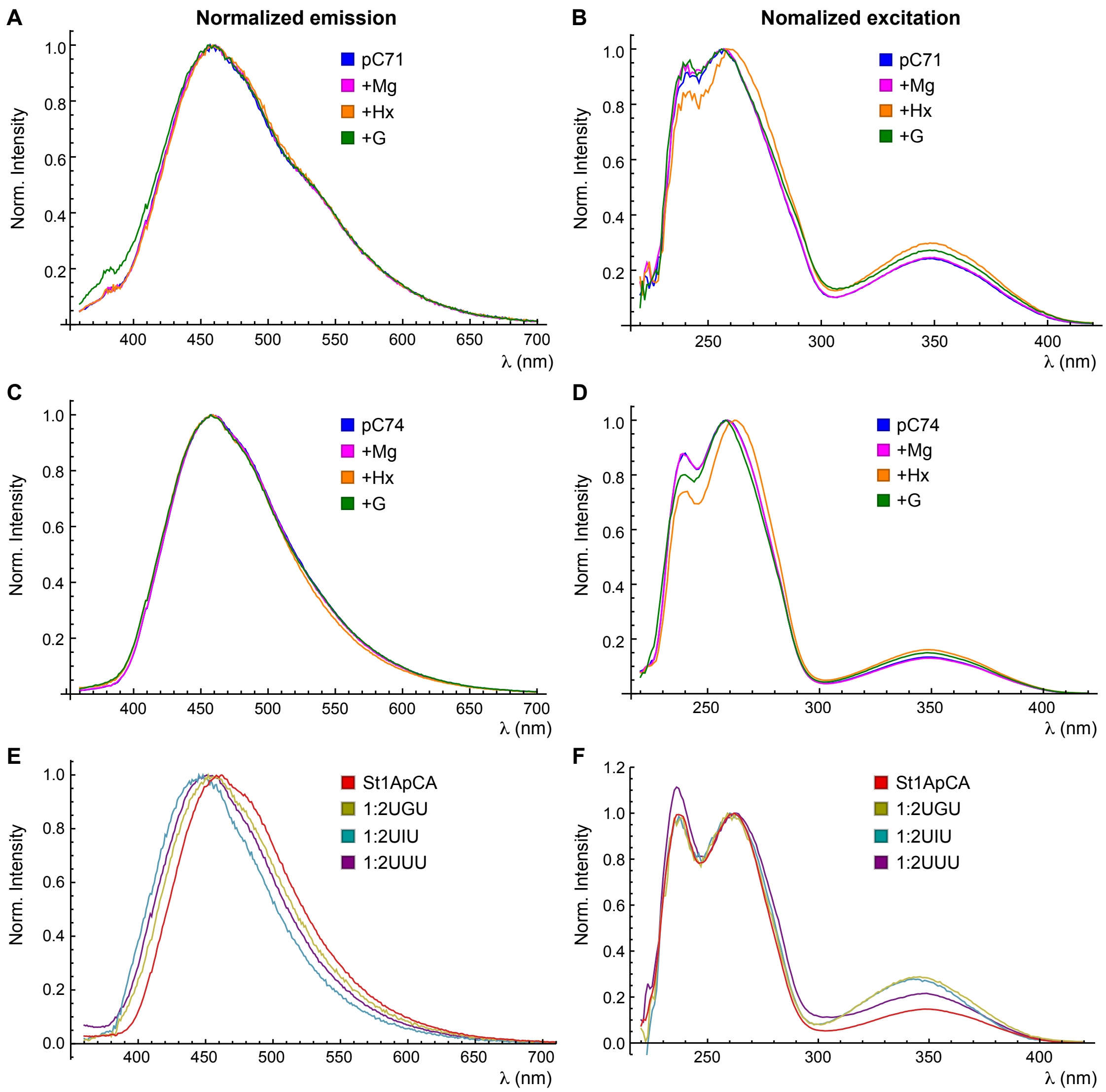


**Figure S2.** Normalized fluorescence emission and excitation spectra. (**A**) Normalized emission spectra reported for pC71 GR in Fig. 2A. (**B**) Normalized excitation spectra reported for pC71 GR in Fig. 2B. (**C**) Normalized emission spectra reported for pC74 GR in Fig. 4A. (**D**) Normalized excitation spectra reported for pC74 GR in Fig. 4B. (**E**) Normalized emission spectra reported for model 21-mers in Fig. 3A. (**F**) Normalized excitation spectra reported for model 21-mers in Fig. 3B.

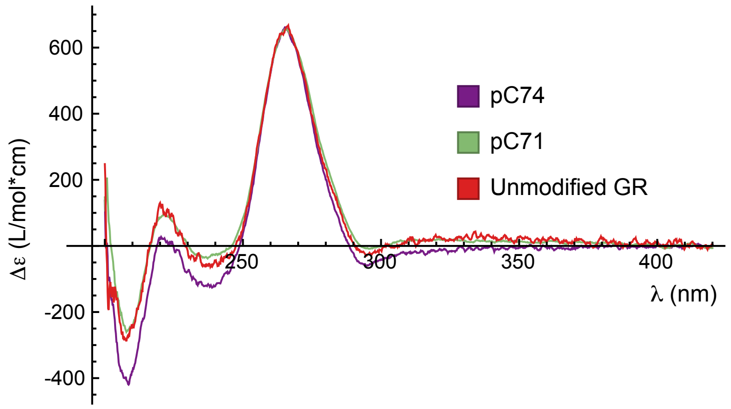


**Figure S3.** CD spectra of pC-labeled and unlabeled GR. Purple: pC74 GR; Green: pC71 GR; red: unmodified GR.

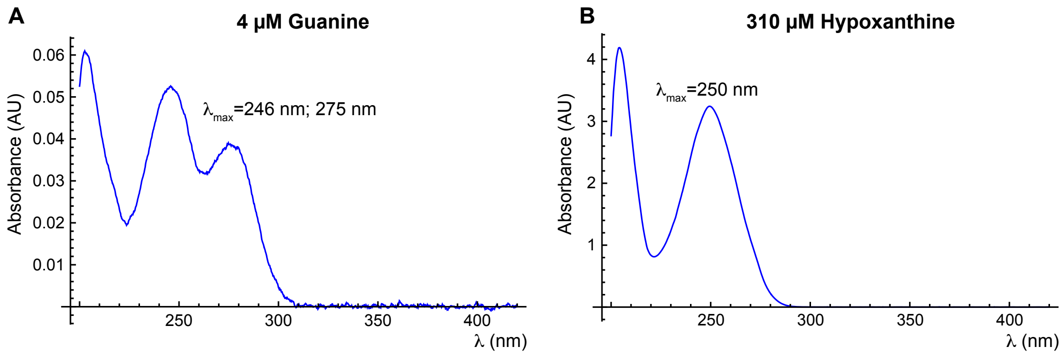


**Figure S4.** Absorbance spectra of G and Hx in a 1 cm pathlength cuvette at the concentrations used in this study. (**A**) Absorbance spectrum of 4 μM G in standard phosphate buffer supplemented with 1 mM $MgCl_2$. (**B**) Absorbance spectrum of 310 μM Hx in standard phosphate buffer supplemented with 1 mM $MgCl_2$.

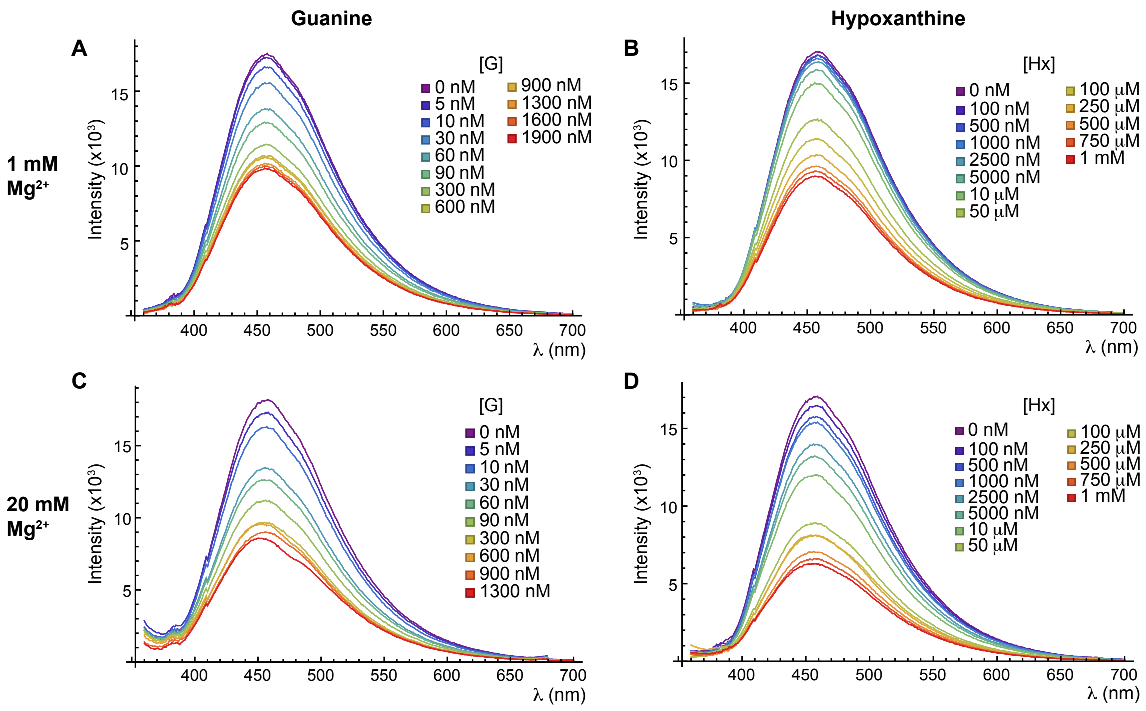


**Figure S5.** Emission spectra of pC74 GR used to construct titration curves in Fig. 5. (**A**) Emission spectra recorded at 1 mM $MgCl_2$ and varying G concentration. (**B**) Emission spectra recorded at 1 mM $MgCl_2$ and varying Hx concentration. (**C**) Emission spectra recorded at 20 mM $MgCl_2$ and varying G concentration. (**D**) Emission spectra recorded at 20 mM $MgCl_2$ and varying Hx concentration.

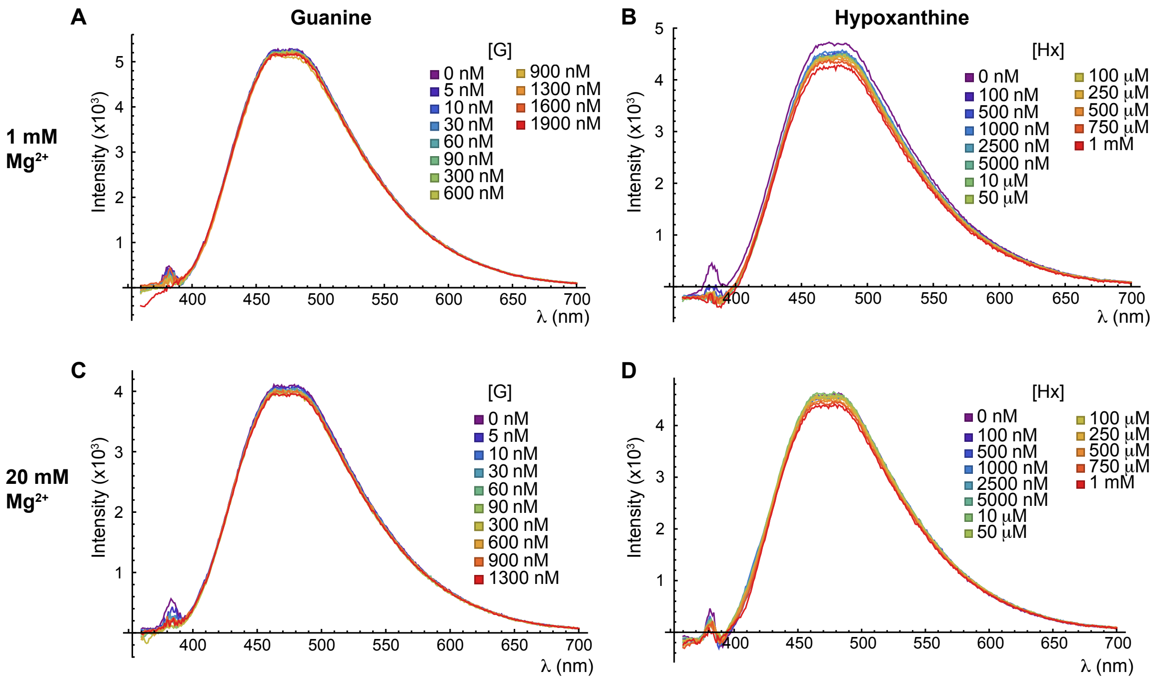


**Figure S6.** Control titrations showing minimal photobleaching or nonspecific response of pC to G or Hx. (**A**) Emission spectra of pCr recorded at 1 mM $MgCl_2$ and varying G concentration. (**B**) Emission spectra or pCr recorded at 1 mM $MgCl_2$ and varying Hx concentration. (**C**) Emission spectra of pCr recorded at 20 mM $MgCl_2$ and varying G concentration. (**D**) Emission spectra of pCr recorded at 20 mM $MgCl_2$ and varying Hx concentration.

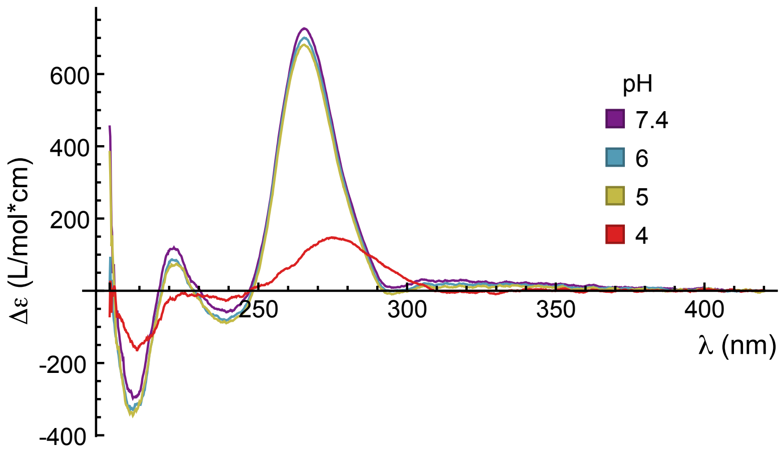


**Figure S7.** CD spectra of unmodified GR at varying pH. The decrease in signal indicates that significant unfolding of helical domains occurs at pH 4.

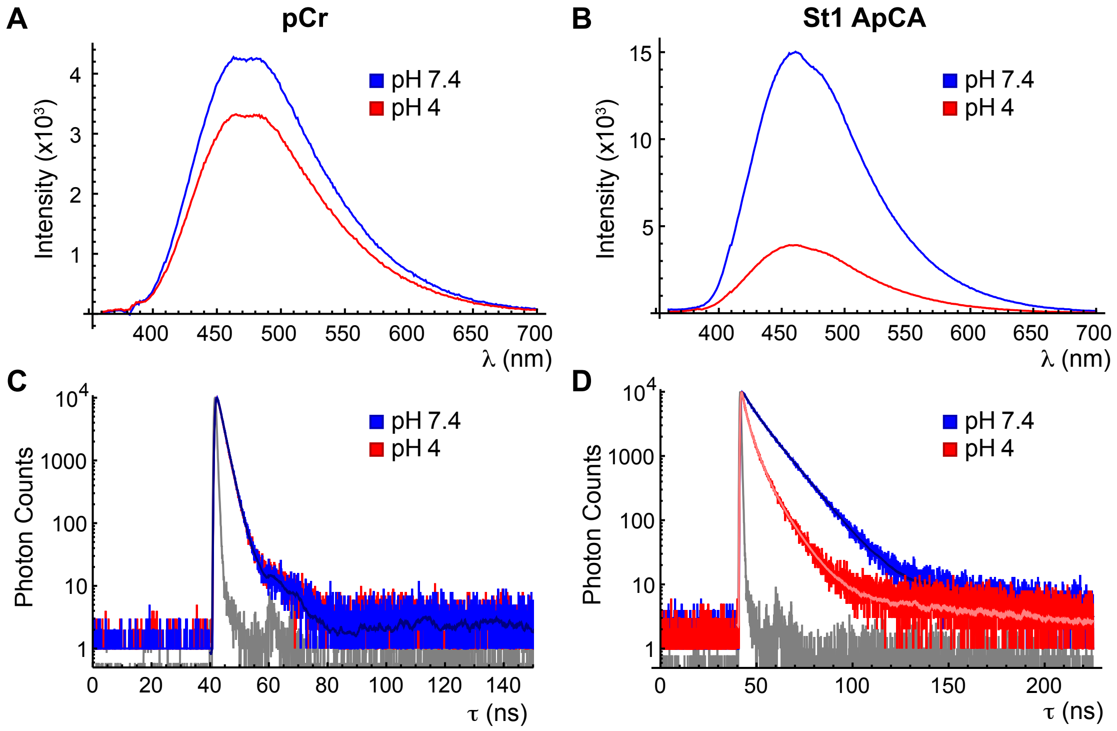


**Figure S8.** Acid denaturation controls. (**A**) Fluorescence emission spectra of pCr in standard phosphate buffer at pH 7.4 (blue) or phosphate buffer adjusted to pH 4 using HCl (red). (**B**) Same as panel **A** for St1ApCA. (**C**) Fluorescence decays of pCr at pH 7.4 (blue) and pH 4 (red). 2E fits are overlaid in contrasting shade. (**D**) Fluorescence decays of St1ApCA at pH 7.4 (blue) and pH 4 (red). 2E (pH 7.4) and 3E (pH 4) fits are overlaid in contrasting shade.

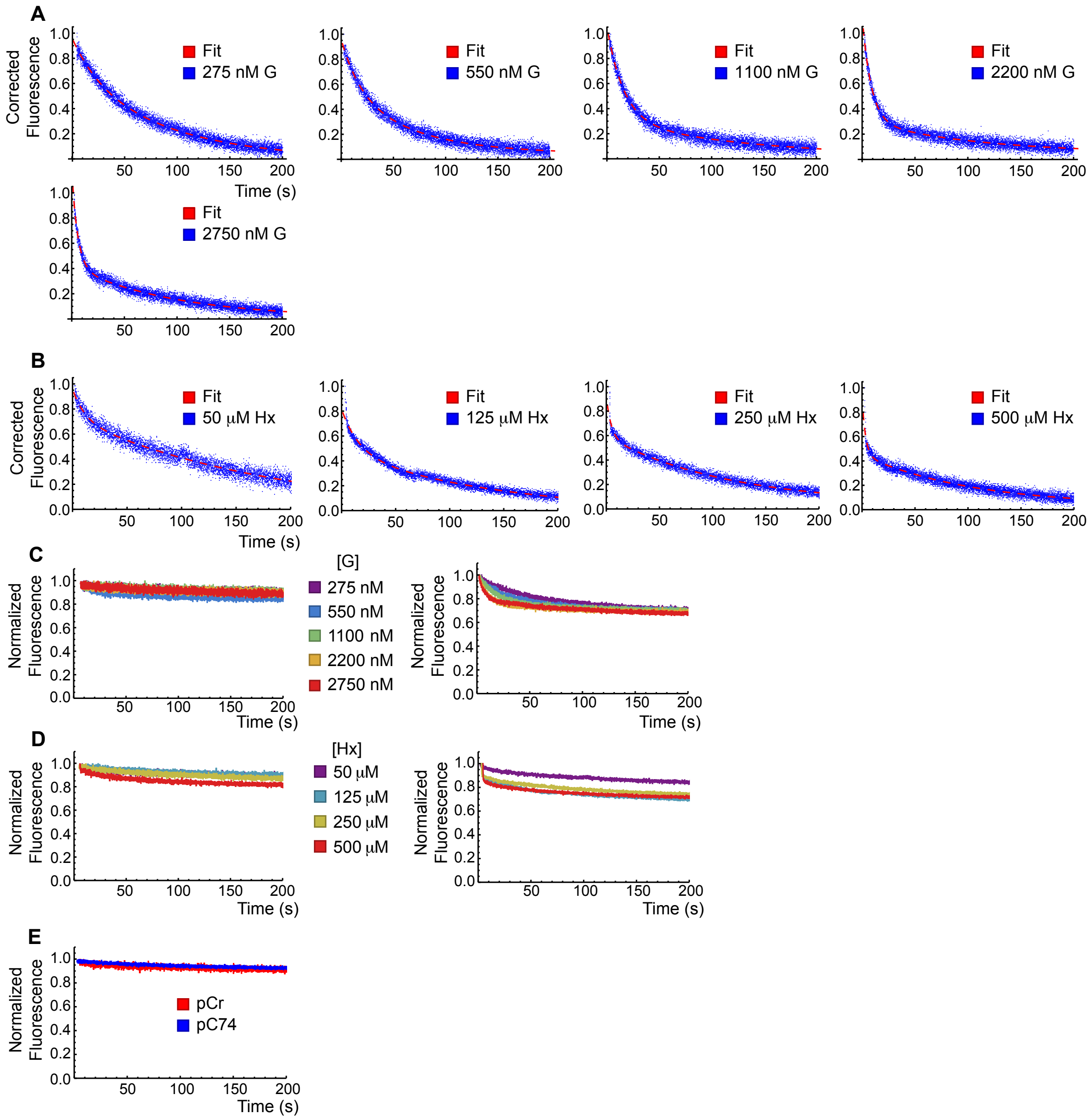


**Figure S9.** Stopped-flow fluorescence spectroscopy fitting and controls. (**A**) Corrected fluorescence traces for association of varying concentrations of G to pC74 GR (blue) and bi-exponential fits (red). (**B**) Corrected fluorescence traces for association of varying concentrations of Hx to pC74 GR (blue) and bi-exponential fits (red). (**C**) Fluorescence traces for pCr (left) and pC74 GR (right) after mixing with concentrations of G matching those in panel **A**. Intensity was normalized to a peak value of 1. (**D**) Fluorescence traces for pCr (left) and pC74 GR (right) after mixing with concentrations of Hx matching those in panel **B**. Intensity was normalized to a peak value of 1. (**E**) Fluorescence traces for pCr (red) or pC74 GR (blue) after mixing with buffer only. Intensity was normalized to a peak value of 1.

**Table S1.** Bi-exponential fitting results for stopped-flow fluorescence spectroscopy. Reported weights are the exponential component amplitudes as a fraction of 1, and may not add up to 1 due to rounding. The uncertainty reported for all rate constants is the standard error resulting from fitting a single fluorescence trace with a bi-exponential function (Equation 8). The error associated with the raw amplitudes was propagated to obtain the uncertainty of the normalized weights displayed below. The errors of rate constants and raw amplitudes were propagated to obtain the uncertainty of the reported $k_{obs}$ value, which is a weighted average of the two rate constants (Equation 9).

| [Ligand] | $R^2$ | $k_{obs}$ ($\bullet 10^{-3}$ $s^{-1}$) | $w_1$ | $k_1$ ($\bullet 10^{-3}$ $s^{-1}$) | $w_2$ | $k_2$ ($\bullet 10^{-3}$ $s^{-1}$) |
|---|---|---|---|---|---|---|
| 275 nM G | 0.98 | 19±2 | 0.30±0.04 | 39±4 | 0.70±0.04 | 10.8±0.8 |
| 550 nM G | 0.97 | 33±2 | 0.43±0.02 | 56±3 | 0.57±0.02 | 15.1±0.8 |
| 1100 nM G | 0.96 | 46.6±0.8 | 0.715±0.004 | 62±1 | 0.285±0.004 | 7.2±0.8 |
| 2200 nM G | 0.96 | 93±1 | 0.761±0.002 | 119±2 | 0.239±0.002 | 9.3±0.5 |
| 2750 nM G | 0.97 | 124±1 | 0.654±0.003 | 184±3 | 0.346±0.003 | 10.2±0.2 |
| 50 μM Hx | 0.96 | 25±1 | 0.233±0.005 | 91±5 | 0.767±0.005 | 5.2±0.1 |
| 125 μM Hx | 0.98 | 29.9±0.7 | 0.414±0.003 | 62.7±0.1 | 0.586±0.003 | 6.8±0.1 |
| 250 μM Hx | 0.97 | 89±4 | 0.38±0.01 | 223±11 | 0.63±0.01 | 8.69±0.007 |
| 500 μM Hx | 0.94 | 168±7 | 0.53±0.02 | 308±15 | 0.47±0.02 | 9.6±0.2 |
